# Brain activity discriminates acoustic simulations of the same environment

**DOI:** 10.1101/2024.08.29.610373

**Authors:** Viola G Matten, Rüdiger Stirnberg, Steven van de Par, Stephan D Ewert, Virginia L Flanagin

## Abstract

In complex acoustic environments, sound localization involves the integration of numerous interrelated auditory and cognitive cues, making it challenging to understand their relationship to brain activity. Here, we use virtual acoustics to probe the brain’s response to auditory distance cues in a realistic environment. We developed a system to record the actual MRI environment, simulated the same room with different degrees of accuracy, then presented sounds at one of two locations in the room. We implemented a novel auditory fMRI sequence to record brain activity. Despite only minor differences in acoustics between the auralizations, it was possible to decode all three rooms from brain activity. A systematic analysis revealed that the direct-to-reverberant energy ratio (DRR) drove brain activity differences between auralizations, centered on the posterior auditory cortex (AC). The results provide strong evidence that the posterior AC processes DRR for spatial auditory perception.

**Impact statement:** A novel fMRI sequence and recording technique are combined with virtual acoustics and multi-variate analyses to decode room simulations from brain activity during distance perception and identify the auditory factors that drive the pattern of activity in the brain.

## Introduction

Auditory perception, unlike vision, is not inherently spatial. The spatial perception of sound requires the conversion of a one-dimensional change in air pressure into a three-dimensional representation of the environment ***Bregman (1990)***.

Nonetheless, hearing provides spatial information about the environment that vision does not capture. For example when an object is placed behind the observer, is visually occluded, or when vision is impaired, hearing can supply information needed to localize objects or navigate through a complex environment. Although we have a good understanding of how and where sound location in azimuth is computed (for reviews see ***Kolarik et al. (2016)***; ***Zahorik et al. (2005)***; ***McAlpine (2005))***, the cues and computations that the brain performs for ADP are much less well understood.

The AC, located bilaterally along the Sylvian fissure, is a crucial region for processing spatial sound. An intact AC is a prerequisite for the ability to localize sound sources ***Thompson and Cortez (1983)***; ***Zatorre and Penhune (2001)***. The posterior part of the AC including the planum temporale and the posterior superior temporal gyrus respond to spatial features of sound ***Rauschecker et al. (1995)***; ***Rauschecker (1997)***; ***Rauschecker and Tian (2000)***; ***Rauschecker (1998)***; ***Ahveninen et al. (2006)***. The posterior AC responds to changes of the sound source in azimuth ***Ahveninen et al. (2006)***; ***Brunetti et al. (2005)***; ***Deouell et al. (2007)***; ***Tata and Ward (2005)***; ***Kopčo et al. (2020)***, distance ***Kopčo et al. (2012)***; ***Kopčo et al. (2020)***, and movement ***Krumbholz et al. (2005)***; ***Warren et al. (2002)***. What auditory spatial information is encoded in the AC is still a matter of debate. It is unclear whether neurons in the non-primary AC are sensitive to basic cues such as interaural level differences (ILD), DRR or intensity or whether they represent higher level features of space such as an abstract representation of sound source location ***Ahveninen et al. (2014)***.

Part of this uncertainty stems from the properties of the auditory cues used for spatial processing, in particular those relevant for ADP. Auditory distance cues are relative in nature, i.e., they rely on additional information, and they differ in their effectiveness ***Kolarik et al. (2016)***; ***Zahorik et al. (2005)***. Distance estimates are more accurate for laterally presented sounds, demonstrating the value of binaural cues ***Kopčo and Shinn-Cunningham (2011)***. However, changes in interaural time differences with distance are below just noticeable differences (JNDs), and ILDs are only modified by variations in distance for distances below 1 m ***Brungart et al. (1999)***; ***Duda and Martens (1998)***. The intensity of the sound, or sound level, is one of the most salient cues for ADP ***Von Békésy (1938)***; ***Gamble (1909)***, however, it is a relative cue. Therefore, the brain cannot determine the distance to a sound source based on sound level alone; knowledge about the initial level of the sound source itself is also required ***Mershon and King (1975)***; ***Zahorik et al. (2005)***; ***Kolarik et al. (2016)***. Reverberation also provides prominant distance information over a wide range of distances, provided the listener has previous information about the environment. While reverberation reduces localization accuracy in azimuth ***Hartmann (1983)***, it improves ADP ***Von Békésy (1938)***. Specifically, the direct-to-reverberant energy ratio (DRR), which is the ratio of the energy of the direct sound to the energy level of the reflections from the walls, provides a reliable distance estimate of sound source distance ***Bronkhorst and Houtgast (1999)***; ***Kopčo and Shinn-Cunningham (2011)***; ***Mershon et al. (1989)***; ***Mershon and King (1975)***; ***Zahorik (2002a,b)***; ***Larsen et al. (2008)***.

The effectiveness of auditory distance cues can also be modified by cognitive factors and other reverberation-related cues. Prior exposure or familiarity with the sound source allows us to better interpret intensity cues, thereby improving distance perception ***Philbeck and Mershon (2002)***; ***Demirkaplan and Hacıhabiboğlu (2020)***; ***Wisniewski et al. (2012)***. When highly familiar sounds are presented, subjects rely less on DRR and more on sound level cues ***Zahorik (2002a)***. Additional properties of the acoustic environment, such as reverberation time (RT30), modify the effectiveness of DRR ***Kolarik et al. (2013)***; ***Schutte (2021)***. Although DRR distance cues, unlike level cues, do not require familiarity with the acoustic environment, prior exposure to the reverberant sound field results in a sustained improvement in the accuracy of distance judgments ***Mershon et al. (1989)***; ***Shinn-Cunningham (2000)***; ***Kopčo et al. (2004)***; ***Zahorik et al. (2005)***. This improvement is stable across different listener positions within the same room ***Kopčo et al. (2004)***; ***Zahorik et al. (2005)***. Likewise, unexpected changes in room reverberation can have a detrimental effect on auditory spatial perception, demonstrating the role of expectation of the surrounding room acoustics on auditory spatial perception ***Brandenburg et al. (2020)***; ***Sutojo et al. (2020)***. Room divergence, for example, is when a room auralization differs from the room subjects are physically located in. Divergence decreases sound externalization ***Werner et al. (2016)***; ***Klein et al. (2017b)***; ***Brandenburg et al. (2020)***, which can be detrimental for auditory spatial perception (for a review see ***Best et al. (2020))***.

The main challenge for understanding the neural underpinnings of auditory distance information is to disentangle the contributions of the different interdependent factors that influence ADP ***Kanwisher (2010)***. Although basic processing and the extraction of binaural cues occurs already at the level of the brainstem ***Grothe et al. (2010)***; ***Jeffress (1948)***; ***McAlpine (2005)***, binaural cues still modulate the neural response in the AC in particular the posterior part of the contralateral superior temporal gyrus ***McLaughlin et al. (2016)***; ***Phillips and Irvine (1981)***; ***Zhang et al. (2004)***; ***Ungan et al. (2001)***; ***Stecker et al. (2015)***; ***Palomäki et al. (2005)***. Similarly, an increase in sound intensity enhances the measured response throughout the whole ascending auditory pathway ***Rapin et al. (1966)***; ***Hegerl et al. (1994)***; ***Neukirch et al. (2002)***; ***Jäncke et al. (1998)***. However, the exact locations that are tuned to intensity are still disputed ***Röhl and Uppenkamp (2012)***; ***Brechmann et al. (2002)***; ***Hart et al. (2003, 2002)***; ***Mulert et al. (2005)***; ***Gutschalk et al. (2002)***. Sound intensity and its perceptual correlate loudness have a complex relationship that further impedes our understanding of how they are processed in the brain ***Röhl and Uppenkamp (2012)***. Activity in the AC is more closely related with the perceived loudness rather than the physical sound intensity ***Langers et al. (2007)***; ***Röhl and Uppenkamp (2012)***, suggesting that the transformation from sound level to loudness is completed at the level of the AC.

In general, a more abstract representation of sound appears to be present in the AC ***Langers et al. (2007)***; ***Röhl and Uppenkamp (2012)***. Whether the posterior AC also represents abstract spatial acoustic concepts is still unknown. Activity in the posterior AC reflects auditory distance, independent of sound intensity cues ***Kopčo et al. (2012)***. This happens also in the frontal plane, where DRR is the predominant distance cue in the absence of intensity cues ***Kopčo et al. (2020)***. These results suggest that the posterior AC is either responding specifically to reverberation cues, or it corresponds to an abstract representation of sound location. Until now the experimental methods and analysis techniques have proved unsuitable to disentangle these abstract concepts from individual auditory cues.

Fortunately, acoustic room simulations provide a technique to manipulate acoustic cues individually by simulating both sound sources and the surrounding environment. Recent advancements in spatial acoustic simulations provide a high level of perceptual plausibility, although a perfect physical reproduction might not be necessary for perceptual plausibility ***Arend et al. (2021)***; ***Hassager et al. (2016)***. With these simulations, it is possible to create physically impossible environments or those that vary in individual acoustic parameters. This opens up the ability to investigate auditory spatial perception in complex, realistic environment in a controlled manner. Instead of manipulating individual parameters like DRR or intensity, we can create a naturalistic experiment by simulating rooms with different acoustic properties and measuring the brain’s response to them.

Concomitantly, advanced multivariate analyses have developed that provide a means to test complex and non-linear relationships between functional magnetic resonance imaging (fMRI) data and the stimuli presented. For example, RSA, allows for a direct comparison between the activity patterns measured with fMRI and computational models describing the processing of sensory stimuli ***Kriegeskorte et al. (2008)***. An abstract representation of the activity maps are computed from the data and then correlated with model predictions based on the stimuli presented and/or behavioral outcomes. With this approach it is possible to quantitatively assess the degree to which each acoustic parameter is represented in the brain response without the need to explicitly define a mapping between the computational models and the imaging data.

Here we used recorded and simulated virtual acoustic environments to explore brain activity during a distance discrimination task, using fMRI. First, we auralized the room where the experiment was conducted, to reduce room divergence and thereby reduce in-head localizations and increase familiarity ***Brandenburg et al. (2020)***. To measure and record the acoustics of the MRI scanner room we built an MRI compatible head and torso model, adapted from a Kemar simulator ***Braren and Fels (2020)*** to capture the binaural impulse response (BRIR). In addition to the measured BRIR, two more BRIRs were generated using room acoustic simulations of the same MRI room with varying degrees of accuracy. Speech sounds were presented via MRI compatible headphones at one of two lateral positions and convolved with one of the three BRIRs. To examine brain activity during distance judgement, subjects were asked to discriminate whether the sound came from the close or far position. An explicit distance task focussed subjects’ attention on the relevant auditory cues, to enhance brain activity in non-primary cortices ***Rämä and Courtney (2005)***; ***Grady et al. (1997)***; ***Jäncke et al. (1998, 1999)***; ***Jancke and Shah (2002)***; ***Petkov et al. (2004)***. Data were collected with a novel MRI scanning sequence that provides ISSS fMRI data acquisition with silent periods for auditory stimulus presentation ***Schwarzbauer et al. (2006)***, but using a 3D-EPI sequence with high spatial and temporal resolution and improved tSNR in subcortical brain regions ***Stirnberg and Stöcker (2021)***. Using a combination of traditional data analysis methods and pattern extraction methods, including multi-voxel pattern analysis (MVPA) and RSA, we found evidence of a separation of room auralization and distance in brain activity. The three room simulations could be decoded from brain activity. A model with the acoustic parameters of each room and distance was compared to brain activity with RSA. It revealed that only DRR contributed to separation of both BRIR and distance. Both the decoding and the RSA analyses were driven by signals in the posterior AC as well as auditory association areas relevant for processing reverberation-related cues. The results suggest the presence of neurons sensitive to DRR in the posterior AC, and demonstrate the usefulness of virtual acoustic simulations and RSA in understanding how we process auditory spatial information.

## Results and discussion

### The reduction of auralization complexity alters distance perception

Acoustic measurements of the MRI magnet room have thus far been inhibited by the high magnetic field and constant background noise generated by cooling systems and helium (re-)compression. However, we developed a system that enabled us to record BRIRs inside the MRI scanner room. The receiver was positioned approximately in the middle of the MRI room on the patient table in front of the MRI ***(Figure 1a)***. Two sound source locations were convolved with one recorded BRIR (*Recorded*) and two simulated BRIRs that capture the acoustics of the MRI magnet room (see Materials and methods for details). The two simulated BRIRs were created using the room acoustical simulator RAZR ***Wendt et al. (2014)***: One simulation modeled the magnet room as closely as possible (*RAZR*), while neglecting objects inside the room. The other simulation was further simplified by auralizing only first order specular reflections and removing the late reverberation, reducing the simulation to a 1st-order image source model (*1st-order ISM*). This led to slight differences between the *Recorded* and *RAZR* BRIRs while the removal of the late reverberation substantially modifies the reverberant part of the impulse response ***(Figure 1b)***.

**Figure 1.**
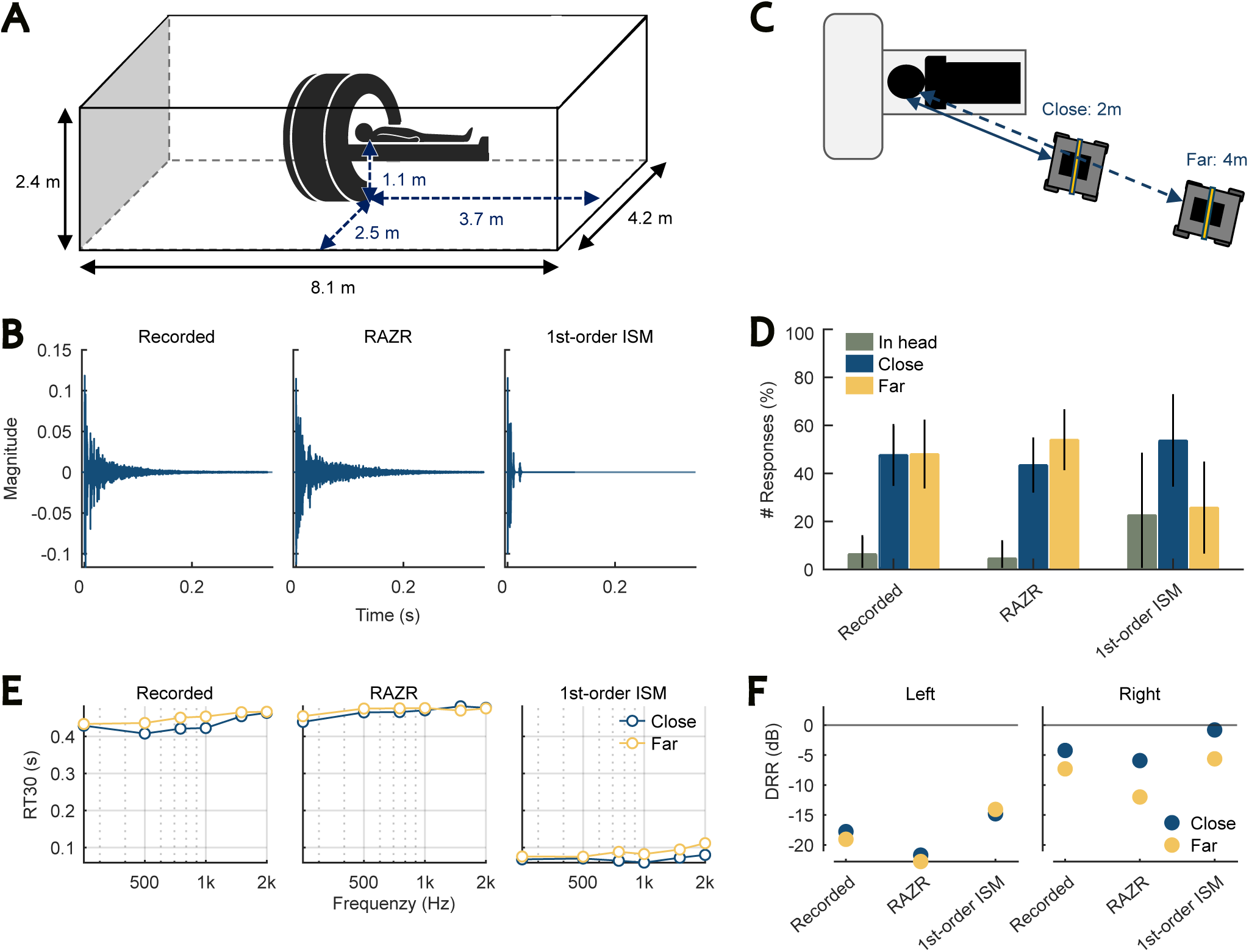
Three sets of BRIRs auralized the MRI room with varying degrees of accuracy. (A) MRI room dimensions. The receiver was located on the patient table in front of the scanner bore, at the meter values shown. (MRI icon: © kilroy79 |Adobe Stock) (B) Representative BRIRs for each auralization. The recorded MRI room (*Recorded*, Left) is compared with two simulations generated with the room acoustic simulator RAZR ***Wendt et al. (2014)***. One simulation modeled the acoustics as accurately as possible (*RAZR*, Middle), the other simulation was a simplified 1st-order ISM without late reverberation (*1st-order ISM*, Right). BRIRs were peak normalized across auralizations. (C) BRIRs were generated at two distances for each auralization. The loudspeakers were placed on a cart to align the vertical position with the subject’s head and approximately 30 degrees to the right of straight ahead. (D) Subjects performed a distance discrimination task between the two distances shown in C. The distribution of responses for each auralization is plotted here. The number of *In head*, *Close* and *Far* responses is normalized by the total number of responses per auralization, resulting in a percent value for each response. Subsequently, the mean percentage value was calculated across all 30 subjects. Error bars indicate the standard deviation from the mean. (E-F) We then performed an auditory analysis of the three room auralization. RT30 and DRR are plotted for the three BRIRs corresponding to one of the distances and auralization type. Labels for the auralizations as in B.

To investigate how the differences in auditory cues influence the neural processing of spatial acoustics, we used a distance discrimination task to focus the subjects’ attention on room acoustical properties. For each auralization, BRIRs were generated at one of two distances ***(Figure 1c)***. The two distances were chosen to maximize the distance difference between the sound sources given the geometry of the magnet room, but close enough to the JNDs to challenge the participants and maintain a high level of attention (see ***Appendix 1)***. Subjects indicated whether they located the sound at their feet (*Close*) or in the corner of the room (*Far*). Additionally, to look for differences in externalization between BRIRs, subjects could alternatively indicate whether sounds appeared inside their head and they were not able to assign a distance to the sounds (*In head*). The experiment was optimized for multivariate analyses, focusing on a large amount of data within a single subject and the use of nonparametric analyses. Due to MRI resources, we recruited subjects to a sample size of 30 (for details see Materials and methods, **subsection** Subjects).

Subjects reported perceived differences in auralizations, but, as expected, struggled to correctly identify the distances as near or far. The percentage correct for discriminating distances was not significantly above chance ***(Figure 1d)***. Note, that we used an alpha value of 0.05 for all statistical tests.

To investigate differences between the response distributions of the three auralizations, we conducted a mulitnomial logistic regression (see Materials and methods). If we compare the *1storder ISM* with *Recorded*, while the distance is held constant, the likelihood of subjects’ responding *In head* was significantly increased compared to the responses *Close* (Wald-*z*(1.35) = 11.034, *p* < 0.001) and *Far* (Wald-*z*(−2.24) = −17.4, *p* < 0.001). Similarly, the comparison between the *1st-order ISM* and *RAZR* revealed that subjects pressed significantly more often *In head* instead of *Close* (Wald-*z*(1.62) = 11.89, *p* < 0.001) or *Far* (Wald-*z*(2.72) = 19.23, *p* < 0.001) in the ISM. Together the amount of *In head* responses was significantly increased in the *1st-order ISM* (more than 20 %) compared to the *Recorded* and the *RAZR* BRIRs (less than 10 %). Hence, subjects were less likely to assign a distance to the sound source when late reverberation was removed. This finding is in line with literature reporting an increase in internalization of sounds when reverberation is reduced ***Hoppe et al. (accessed 27 September 2024)***; ***Begault et al. (2001)***; ***Catic et al. (2015)***; ***Leclère et al. (2019)***; ***Toole (1970)***; ***Best et al. (2020)***. In contrast, internalization did not differ between *Recorded* and *RAZR*. A binomial logistic regression comparing the number of *In head* responses with the sum of *Close* and *Far* responses revealed no significant difference. We conclude that compared to the recorded BRIRs the full RAZR simulation achieves a similar spatialisation, i.e. the impression that the sound source is located in a three-dimensional environment.

Moreover, for the *1st-order ISM*, the likelihood of subjects’ responding *Close* instead of *Far* is significantly increased compared to *RAZR* (Wald-*z*(1.09) = 15.48, *p* < 0.001) and *Recorded* (Wald-*z*(0.89) = 12.52, *p* < 0.001). Therefore, sounds auralized with the *1st-order ISM* are perceived closer compared to the two BRIRs with enhanced reverberation, consistent with literature relating a reduction in reverberation to smaller distance estimates ***Butler et al. (1980)***; ***Mershon et al. (1989)***; ***Zahorik et al. (2005)***; ***Kolarik et al. (2016)***.

To quantitatively assess acoustic differences in the auralizations with the three different sets of BRIRs, an auditory analysis was performed that extracted room-acoustical parameters known to be important for characterizing the perception of room acoustics. Among others, these parameters include loudness, ILD, RT30 and DRR (see ***Appendix 1)*** for all auditory parameters measured). The perceptible acoustic differences in the three auralizations (*Recorded, RAZR*, or *1st-order ISM*) should be related to these acoustic parameters. The auditory analysis revealed that the auralizations were highly similar, specifically between the *Recorded* and the realistic simulation *RAZR*. Primary differences were seen for the reduced simulation *1st-order ISM*, although differences could be found between all three auralizations. For example, RT30 and DRR are plotted here ***(Figure 1e*** and ***Figure 1f***, for all parameters see ***Appendix 2 — Figure 1)***. Removing the late reverberations resulted in a reduction in RT30 ***(Figure 1e)***. The simultaneous increase in *In head* responses is consistent with findings in the literature relating a decrease in RT30 with an increase in internalization ***Catic et al. (2013)***; ***Begault et al. (2001)***; ***Toole (1970)***.

Although the degree of internalization is similar between the *Recorded* and the *RAZR* auralizations, distance perception differs between the two. The ratio between *Far* and *Close* was significantly increased for *RAZR* compared to *Recorded* (Wald-*z*(0.21) = 3.26, *p* = 0.001). In addition, we find an increase in the likelihood of subjects responding *Far* instead of *In head* for *RAZR* compared to *Recorded* (Wald-*z*(0.48) = 3.08, *p* = 0.002). As shown in ***Figure 1d*** we observed more *Far* responses than *Close* for *RAZR* while for *Recorded* the ratio between *Close* and *Far* was equal. However, note that the differences between *RAZR* and *Recorded* are much smaller than for the comparison with the *1st-order ISM*. The observation is in line with a recent study showing that the two perceptual quantities: distance and internalization, are differentially affected by changes in audio rendering and reverberation ***Hoppe et al. (accessed 27 September 2024)***. The results suggest that distance perception is slightly enhanced if sounds are auralized with RAZR instead of the recorded BRIRs. This may be related to the acoustic effects of the fMRI that are unfamiliar to most observers and were omitted in the *RAZR* auralizations.

The DRR values would predict a similar variation in distance perception ***(Figure 1f)***. DRR is slightly reduced for *RAZR* compared to the *Recorded* BRIRs, which is known to increase perceived distance ***Zahorik et al. (2005)***; ***Kopčo et al. (2012)***. DRR is strongly increased for the *1st-order ISM*, which should lead to shorter perceived distances. The increase in DRR for the *1st-order ISM* is related to the removal of the late reverberation while the difference between *RAZR* and *Recorded* is likely caused by additional early reflections from the scanner bore during the recording as well as shielding effects of the scanner (for a more detailed description see ***Appendix 1)***. Overall, the logistic regression of the behavioral data demonstrates that subjects could perceive differences between the auralizations, and that the subtle changes in acoustic cues caused by different levels of accuracy in the BRIRs are sufficient to modify ADP.

### Room auralizations recruit the same set of brain regions

Room auralization had a significant effect on perception, in particular the room simulation with less reverberation reduced perceived distance. We then investigated how these differences between auralizations affected brain activity. We conducted a general linear model (GLM) analysis and compared each room auralization to a silent baseline condition (see Materials and methods). We observed activity in primary and nonprimary AC as well as the thalamus and premotor areas. A visualization of the brain areas that were activated can be seen in ***Figure 2a***.

**Figure 2.**
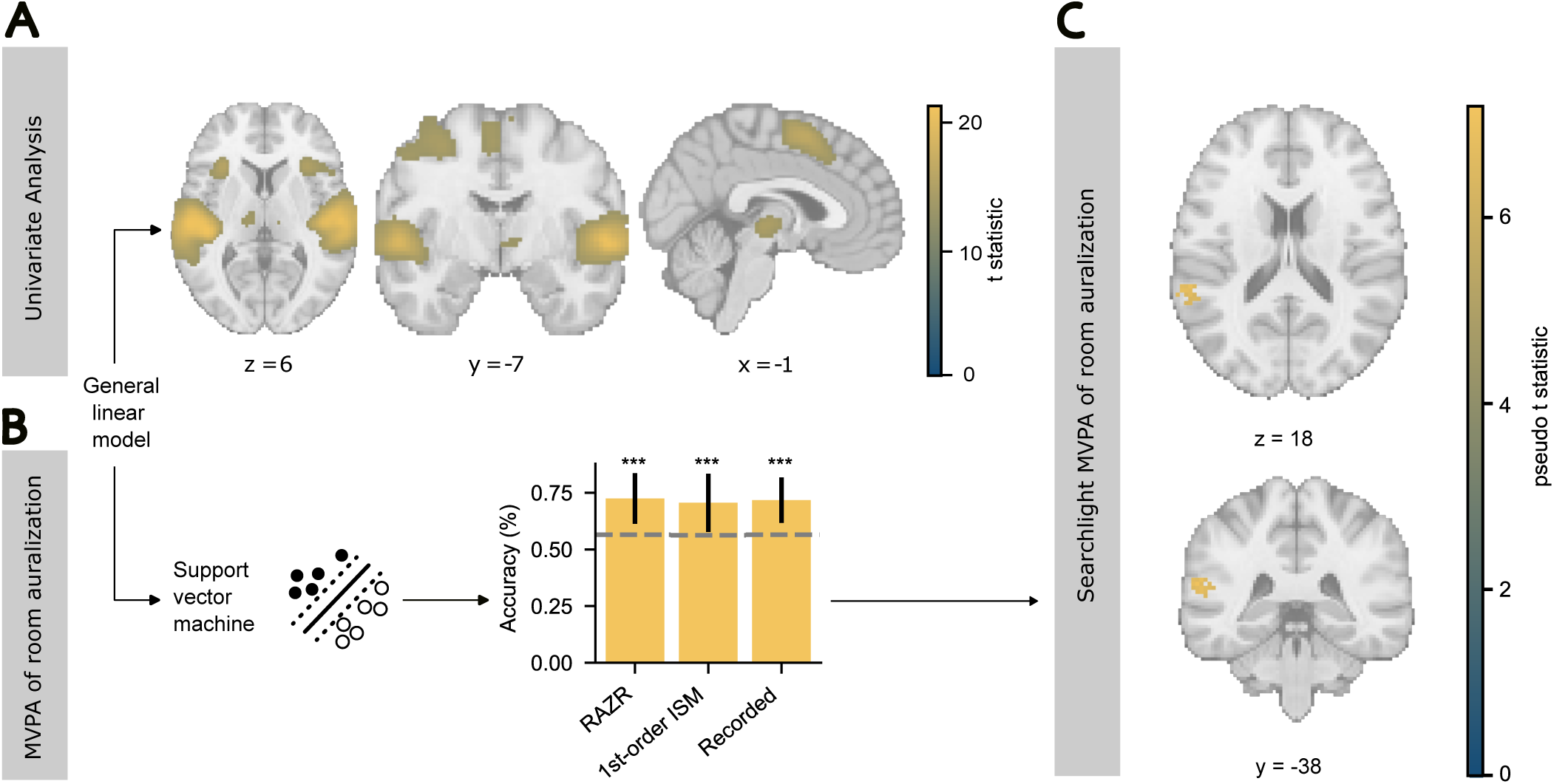
Univariate fMRI analyses showed similarities between room auralizations, whereas multivariate analyses revealed distinct patterns of activation for each room auralization. (A) Univariate regression analysis (GLM) based on the data for the 30 subjects. We observed activity in the auditory regions including AC, thalamus, Heschl’s gyrus and planum temporale. Smaller clusters were found in frontal and motor areas including the insula and the cingulate cortex (see ***Appendix 3 — Table 1*** for details). The statistical map has been thresholded at *p* = 0.05 FWE. Only cluster with more than 100 voxels are considered significant. (B) Multivoxel pattern analysis (MVPA). Decoding accuracy of room auralizations from brain activity. We computed the mean decoding accuracy across all 30 subjects. Error bars show the standard deviation from the mean and dashed lines indicate chance accuracy. For each auralization the average chance accuracy was significantly larger than chance level (*** *p* < 0.001, one-sided t-test). (C) Localization of MVPA. Statistical map of the searchlight based on SVM classifiers distinguishing each BRIR auralization from the other two for each of the 30 subjects. A significant cluster was observed in the left superior temporal lobe. Maps were threshold at *p* = 0.05 FWE and only clusters with more than 50 voxels were considered significant. For further details regarding cluster volume and peak statistic refer to ***Appendix 4 — Table 1***. All brain activity patterns (including (A)) are overlaid on the MNI152 single subject template brain and coordinates are given in MNI space.

The strong activation of the AC and thalamus demonstrates the effectiveness of the imaging sequence. The main challenge for acoustic experiments in the MRI machine is the noise resulting primarily from rapidly alternating readout gradients. The higher the spatial and temporal resolution, the higher the gradient amplitudes and the longer the ramp times, which typically increases the noise. Multiple approaches exist to reduce noise while measuring brain activity. We implemented an EPI sequence with ISSS imaging that alternates loud (with readout gradients) and silent periods (without readout gradients) for stimulus presentation ***Schwarzbauer et al. (2006)***. This approach improves imaging of ascending auditory pathway ***Dewey et al. (2021)***. We extended the method to a multi-shot 3D-EPI sequence, increasing temporal resolution to 0.57 s while maintaining a relatively high spatial resolution of 2.5 mm isotropic and, at the same time, further minimizing acoustic noise (see Materials and methods).

When looking at the brain activity evoked by each auralization individually, neither recruited brain regions nor overall activation strength differed between the auralizations. Therefore, the sounds presented during the distance discrimination task recruit the same set of auditory brain regions independent of auralization. As expected, it was not possible with a classical fMRI analysis to separate out the differences between the room auralizations.

### Room auralization can be decoded from brain activity

Although we observed differences in behavior for the three room auralizations, this was not reflected in the univariate regression analysis (GLM) of brain activity. In other words, differences between the room auralizations did not change the average activation strength of a single brain area, nor did they activate additional brain areas. While a univariate approach answers questions about the size and strength of brain activity, it may not reveal changes in pattern within these regions ***Norman et al. (2006)***; ***Haynes (2006)***. Since the auditory simulations *RAZR* and *1st-order ISM* modeled the *Recorded* room, they differ only slightly in their auditory cues, with values close to JNDs. Therefore, brain activity is most likely modified in a fine-grained fashion by these subtle difference in room auralizations.

We hypothesized that auralizations alter the pattern of brain activity within regions rather than the activation strength. In contrast to the GLM approach, MVPA, or decoding, allows us to identify whether individual conditions evoke a consistent nonlinear change in the brain’s activity pattern ***Norman et al. (2006)***; ***Haynes (2006)***. We therefore used MVPA to test whether room auralizations can be discriminated from brain activity. Three linear support vector machine classifiers were trained on the activity of the entire brain to test whether each room auralizations can be decoded from the other two based on the imaging data. From the eight fMRI runs, we conducted a leave-one-run-out crossvalidation to train the classifier. We were not interested in finding a classifier that can generalize across all subjects, but rather wanted to obtain an estimate of the classification performance for each subject individually. Hence, we did not tune the hyperparameters on an extra validation set to minimize the effect of noise, but instead averaged the crossvalidation scores to obtain an estimate of the accuracy with which the room auralization can be decoded for each subject. Subsequently, accuracies were averaged across subjects.

The final mean accuracy, with which each trained classifier could discriminate one auralization from the other two, can be seen in ***Figure 2b***. Chance accuracy was determined with permutation testing. For each auralization, the classification accuracy was significantly above chance (*Recorded*: t(0.72)=8.29, *p* < 0.001; *RAZR*: t(0.72)=7.80, *p* < 0.001; *1st-order ISM*: *t*(0.71) = 6.00, *p* < 0.001;). There was no significant difference between the mean classification accuracies (*F* (2, 27) = 0.21, *p* = 0.81). Hence, the classifiers performed equally well for all three auralization types. These results confirm our hypothesis that the small differences between the room auralizations could still modify the pattern of brain activation. It is worth noting that we observe this effect although the acoustic parameters varied in other dimensions unrelated to room auralization. The distances as well as the different spoken words introduced variability in the presented sounds that were as large as, or larger than the variations introduced by the different auralization types (see ***Appendix 1*** for the acoustic analysis across distances and ***Stevens (1972)*** for acoustic differences in speech).

### The posterior superior temporal lobe encodes room auralization

After determining that the rooms auralizations could be decoded based on the pattern of activity across the entire brain, we wished to identify the specific brain regions contribute the most to this classification. To achieve this we conducted a searchlight analysis based on a classifier that utilized linear support vector machines with a one-versus-rest multiclass strategy. To assess significance at the group level, we performed a permutation test and determined significance with a variance-smoothed t-statistic to mitigate noise (for more information see ***SnP (2004))***. Additionally, clusters with less than 50 contiguous voxels were considered noise. A cluster with significant decoding accuracy was observed in the left superior temporal lobe including the superior temporal gyrus, the planum temporal and the Heschl’s gyrus ***(Figure 2c)***.

This outcome confirms that the pattern of activity in the superior temporal lobe contains enough information to distinguish between room auralizations. This could be due to the differences in auditory parameters between the room auralizations, or it could reflect and abstract cortical representation of the auditory environment (for a review see ***Ahveninen et al. (2014))***. The posterior AC is known to processes spatial information, but it is still unclear how this information is encoded. Individual neurons may represent specific locations in space or respond to abstract spatial characteristics such as egocentric distance, azimuth or elevation instead of being sensitive to basic acoustic cues.

Recent literature however, has proposed the existence of neuron populations responsive to DRR in the posterior part of Heschl’s gyri, the planum temporale and the superior temporal gyrus ***Kopčo et al. (2012)***; ***Kopčo et al. (2020)***. As the room auralizations here differ primarily in their reverberation (see ***Appendix 1)***, our findings are consistent with this proposal, suggesting a higher sensitivity in the superior temporal lobe to changes in reverberation. Nevertheless, because we were interested in using realistic stimuli, the three room auralizations differed also in other acoustic parameters. The question remained, therefore, what features contributed to the discrimination of the three room auralizations.

### Acoustic parameters for the room auralizations show similarity with brain activity

The classification analysis revealed that changes in the auditory reproduction of the MRI room shape the neural response in a unique and consistent way. To understand the relation between the observed changes in the neural response and the underlying acoustic parameters, we conducted a RSA based on behavior, categorization of the stimulus type and the acoustic parameters of the sounds. While MVPA tells us whether discriminant information about individual stimulus conditions can be found in the multivariate pattern of brain activity, RSA allows us to relate this activity pattern to specific computational models of the stimuli and resulting behavior ***Kriegeskorte et al. (2008)***; ***Xu et al. (2021)***. RSA compares the similarity between activity patterns for different conditions with the similarity predicted by models of the stimulus conditions. Thus, the method enables us to characterize what information is represented in the brain and to correlate it quantitatively with model predictions.

We quantified acoustic parameters that are known to be relevant for perception of spatial acoustics in particular for ADP. These parameters included DRR, RT30, the Mel-Frequency-Cepstral Coefficients (MFCC), interaural cross-correlation (IACC), perceived loudness and ILD ***Kolarik et al. (2016)***; ***Zahorik et al. (2005)***; ***Coleman (1963)***. For DRR, we determined the mean value of the left and the right channel (*mean DRR*) as well as the difference in DRR between the two channels (Δ *DRR*). While *mean DRR* captures the monaural variations in DRR, Δ *DRR* describes the binaural changes in DRR. For each condition this set of parameters was computed. Using basic measures such as pearson correlation or euclidean distance, we determined the dissimilarity between conditions for each parameter. This procedure results in a dissimilarity matrix for each parameter. At the same time, we compared the activity maps obtained from the GLM for each condition resulting in one dissimilarity matrix per subject. Subsequently, the matrices based on the imaging data were averaged across subjects.

We ended up with one dissimilarity matrix that characterized the information in the activity patterns, and a set of dissimilarity matrices describing the acoustics of the presented stimuli. By comparing the matrix based on the data and the matrices based on the acoustic parameters, it was possible to investigate to what degree the observed brain activity contains information related to the various parameters without drawing detailed assumptions about how the parameters might be computed by the brain (see Materials and methods and ***Kriegeskorte et al. (2008))***. As a result, we obtained a correlation coefficient for each acoustic parameter. This coefficient can be interpreted as a prediction accuracy that indicates to what degree each parameter is represented in the brain.

In addition, we created three dissimilarity matrices, one for each response type, that captured the behavioral response. For each condition the relative amount of *In head*, *Close* and *Far* responses was computed. The pairwise dissimilarity of the resulting values was determined using euclidean distance. Furthermore, we included a set of matrices characterizing the category of sounds. Two conditions were defined to have a minimal dissimilarity of 0 if they belonged to the same category (*Recorded*, *RAZR*, or *1st-order ISM*) and 1 otherwise. The different categories were based on the three room auralizations and the distance of the sound source. Incorporating these models allowed us to compare the extent to which the observed brain activity reflects higher-level information in contrast to the basic acoustic parameters. The response type based models indicate the extent to which behavioral choice is represented in brain activity. The categorical models quantify the relationship between brain activity and processes related to the determination of abstract cognitive categories, such as room auralization and distance.

Neither the behavioral nor the categorical dissimilarity matrices were notably correlated with the imaging data ***(Appendix 5 — Table 1)***. The dissimilarity matrices corresponding to the acoustic parameters, however, did show a relationship with the imaging data. The highest correlation i.e. accuracy was observed for the parameters based on DRR ***(Figure 3a)***, demonstrating the importance of this cue. However, the accuracy of the DRR models was not significantly higher than the other individual models. Testing against zero yields p-values above 0.1 for all single acoustic models. Still, this is not surprising, since a single acoustic parameter is unlikely to significantly predict activity across the entire brain.

**Figure 3.**
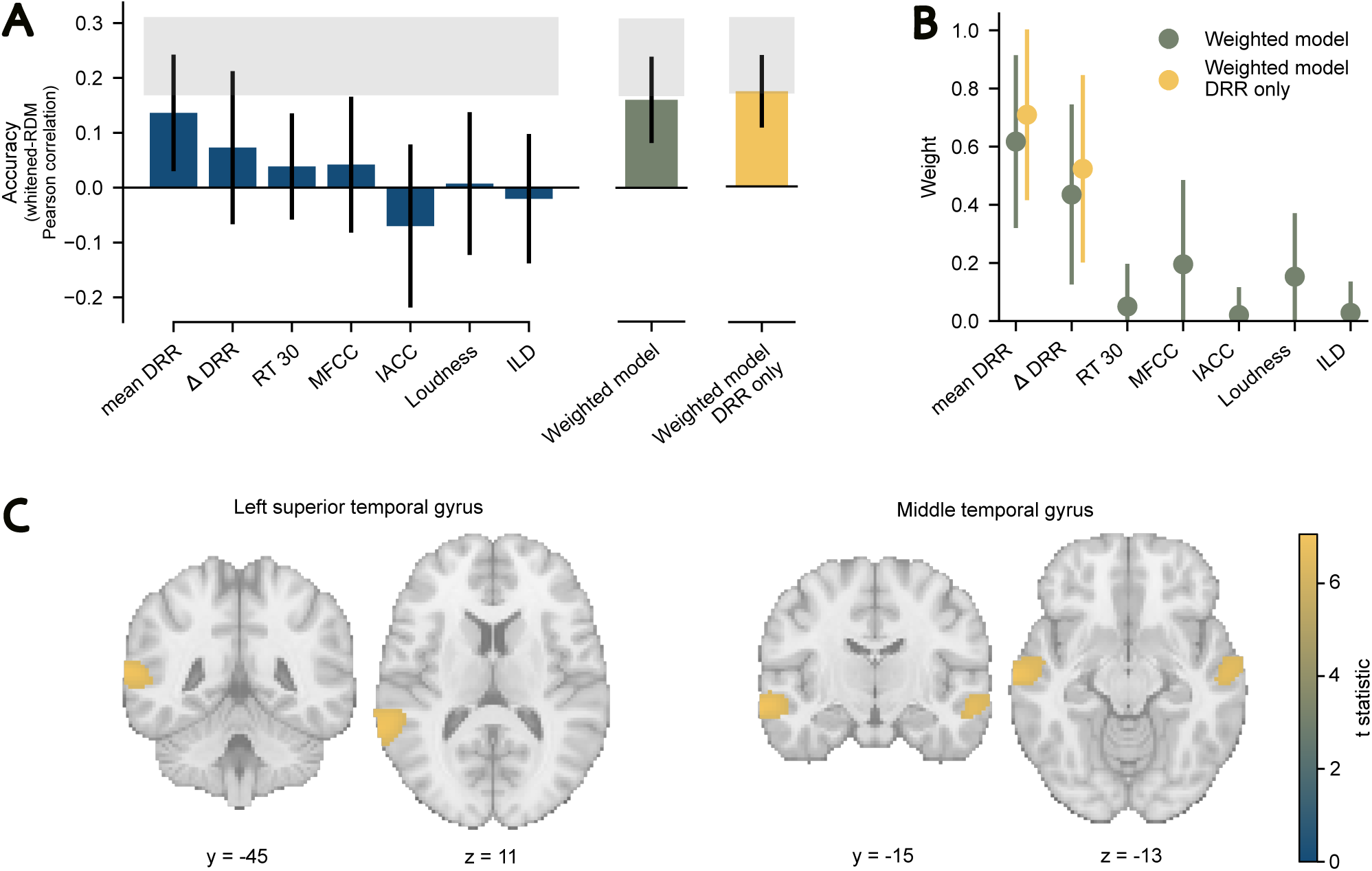
RSA revealed DRR drives brain activity and is predominantly represented in the posterior AC. (A) Prediction accuracy of the models tested based on auditory cues. Error bars were obtained using bootstrapping with *N* = 1000. The gray bar indicates the noise ceiling. It provides an upper and lower bound on the performance of an optimal model given the noise in the data. Blue: Accuracy of models based on individual acoustic parameters. The two DRR parameters differ in the way the left and right channel were combined. *mean DRR* was based in the average of both channels and Δ*DRR* was based on the difference. Green and yellow: Models based on a linear weighted combination of acoustic parameters. *Weighted model* combined all parameters shown on the left. *Weighted model DRR only* combined the two DRR models. (B) The weighted models were fit using crossvalidation and error size was estimated using bootstrapping (*N* = 1000). The plot shows the obtained weights for the weighted model that combined all examined parameters (green) and for the best fitting model containing only the DRR parameters (yellow). (C) Results of searchlight RSA for the *Weighted model DRR only*. A cluster radius of 5 voxels has been used. Images are thresholded at 0.05 FWE and only clusters with more than 50 voxels were considered significant. Brain activity patterns are overlaid on the MNI152 single subject template brain and coordinates are given in MNI space. For details regarding cluster location, size and peak statistic refer to ***Appendix 6 — Table 1***. Data from all 30 subjects were included in the analyses for each plot in this figure.

With the RSA method it is also possible to fit a linear weighted combination of the individual acoustic parameters. This weighted model allows us to evaluate whether brain activity can be related to a combination of parameters or categories and it quantifies the contribution of each parameter to the final fit. Although, the exact combination is presumably highly nonlinear, this approach allows to investigate which parameters are reflected in brain activity and to assess the relative importance of each parameter to the optimal model given the brain activity in the current experiment ***Kriegeskorte et al. (2008)***; ***Xu et al. (2021)***. We therefore fit a weighted model with all of the parameters to the data using crossvalidation. As expected, the full weighted model performed better than the individual models ***(Figure 3a***, Uncorrected, one-tailed t-test against 0: *t*(0.16) = 2.04; *p* = 0.05; see also ***Appendix 5 — Table 1)*** and the weights obtained for the individual acoustic parameters are depicted in ***Figure 3b***. The largest weights were observed for the two DRR models (Table ***Table 1)***. The next highest weight was for loudness, but all of the weights of the remaining parameters ranged from zero to 0.2. The absolute values of the weights do not have a direct interpretation. However, if compared with each other they provide a measure to quantify the contribution of each parameter to the weighted model.

**Table 1.**
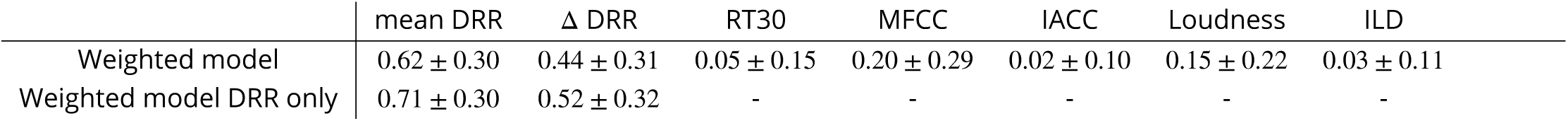
The weights for the full weighted model with all measured acoustic parameters as well as the final weighted model based only on DRR.

| | mean DRR | $\Delta$ DRR | RT30 | MFCC | IACC | Loudness | ILD |
| --- | --- | --- | --- | --- | --- | --- | --- |
| Weighted model | $0.62 \pm 0.30$ | $0.44 \pm 0.31$ | $0.05 \pm 0.15$ | $0.20 \pm 0.29$ | $0.02 \pm 0.10$ | $0.15 \pm 0.22$ | $0.03 \pm 0.11$ |
| Weighted model DRR only | $0.71 \pm 0.30$ | $0.52 \pm 0.32$ | - | - | - | - | - |

Due to the low weights of many of the acoustic parameters in the full model, it was likely that a model with a reduced number of acoustic parameters would fit the brain activity better than the full model. If a reduced model provides a better fit to the imaging data, it would be strong evidence that only certain acoustic parameters were relevant for brain activity during this task. To identify which acoustic parameters improved the fit, we systematically removed individual acoustic parameters from the weighted model and recalculated the accuracy for the remaining model. This approach yielded that the best fitting model was a combination of the two DRR-related parameters (Uncorrected, one-tailed t-test against 0: *t*(0.17) = 2.62, *p* = 0.02). Hence, we conclude that brain activity is best described by a model that captures both monaural and binaural variations in DRR. The observation is in line with the differences in behavior found, suggesting that reverberation is the main cue used by subjects throughout the experiment. These findings reveal that subtle variations in auditory cues are nonetheless represented in brain activity, and that a multivariate approach relating an acoustic analysis of the stimuli to the brain activity can disentangle what parameter are relevant to a task.

Interestingly, although loudness is a strong cue for ADP, and although it had the third highest weight in the full weighted model, leaving it out of the model improved the fit, suggesting that it did not contribute to brain activity in this task. We believe this is primarily a result of the small differences between the two distances, and the lack of an absolute intensity reference, rendering intensity differences a less interpretable distance cue in this task.

### DRR is represented in the posterior auditory and association cortices

In the previous RSA, the neural response of the entire brain was used for the analysis. We assume, however, that the observed correlation between the brain activity and DRR, can primarily be attributed to a set of regions in the AC. To identify whether the intrinsic representation of a certain region resembles the representation predicted by DRR, we conducted a searchlight analysis using the reduced weighted-DRR model. This resulted in a statistical map displaying, for each voxel, the similarity between the model and the activity in the proximity of that voxel ***Kriegeskorte et al. (2006***, 2008).

We observed a high correlation with the DRR model in the posterior AC and the auditory association area including the superior temporal lobe contralateral to the stimulus and the medial temporal lobe ***(Figure 3c)***. Although the remaining auditory cues, such as ILD or RT30, were not significantly represented in overall brain activity, localized activation may still correlate significantly with individual parameters. Therefore, we performed a searchlight RSA for each of the dissimilarity matrices calulated for the whole brain analysis. No significant correlation was found between any of the individual dissimilarity matrices and a specific region of the brain. We hypothesize that subjects focused their attention on DRR as a salient auditory distance cue, rather than on the remaining parameters thereby enhancing the sensitivity to changes in DRR ***Gandhi et al. (1999)***; ***Somers et al. (1999)***; ***Foster and Ling (2022)***. These results suggest that neuronal populations in these areas are sensitive to variations in DRR resulting from changes in distance and BRIR.

The result is consistent with previous work showing that dorsal auditory pathways process auditory location information. Neurons in the posterior non-primary AC encode spatial information about the sound such as horizontal direction changes. ***Rauschecker et al. (1995)***; ***Rauschecker (1997)***; ***Rauschecker and Tian (2000)***; ***Rauschecker (1998)***; ***Ahveninen et al. (2006***,?). Furthermore, the superior temporal gyrus has been associated with reverberation ***Kopčo et al. (2012)***; ***Kopčo et al. (2020)***. Kopčo et al. conducted two adaption fMRI experiments where they varied distance or intensity independently while subjects performed an unrelated task. They found adaptation to distance, independent of intensity and binaural cues, in the posterior AC. Since DRR is the most important remaining cue, they suggested a relationship between reverberation based cues and the posterior AC. However, they could not disentangle whether the observed activity was due to the region encoding an abstract representation of auditory distance or whether it can be attributed to neuronal populations sensitive to reverberation-related cues. Our data provide strong evidence that these areas are specifically sensitive to DRR. Recently, non-primary auditory cortices, including the posterior superior temporal cortex, were demonstrated to be invariant to their responses to sounds with or without background noise ***Kell and McDermott (2019)***. This invariance to a background signal would be a necessary processing step to be able to extract distance relevant information from a soung in a reverberant environment. Taken together, these findings support the hypothesis that neural populations in the posterior AC exist that respond specifically to changes in DRR.

The correlation we found with DRR in the superior temporal lobe was contralateral to the presented stimuli, consistent with the results by Kopčo et al.. However, as sounds were presented only on one side, additional studies are required to verify whether DRR is encoded stronger by the contralateral hemisphere. It is important to note that the observed relationship between DRR and the pattern of activity in the AC could also be caused by additional auditory cues that correlate with DRR. A number of spectral and temporal cues have been suggested to be relevant for the perception of DRR, such as IACC, spectral variance or envelope, as well as buildup and decay times ***Larsen et al. (2008)***; ***Bronkhorst and Houtgast (1999)***; ***Bronkhorst (2001)***; ***Kim et al. (2001)***; ***Kolarik et al. (2016)***. These cues covary monotonically with DRR, and would therefore be difficult to separate out in the present study. We included those cues known to be related to distance perception allowing us to identify the type of distance-relevant acoustic cue represented in brain activity. A next step could be to specifically manipulate the auditory cues that correlate with DRR, which would be possible with the virtual acoustic simulation methods used here. Additionally, semantic perception is also known to activate regions in posterior AC, which could influence the brain activity we found. Although we used speech in all our conditions, it was irrelevant for the task. Also, we counterbalanced the words across distances and room auralizations, so the semantic content was the same across the conditions we tested. Therefore, it is unlikely that our observations are purely a result of semantic processing. Nevertheless, future studies should consider using non-verbal sounds.

We were surprised that the binaural cues such as ILD did not have a strong representation in the brain activity during our task. The sounds came from the right side, and differences in binaural cues for the different distances and BRIRs were seen in the auditory analysis (see ***Appendix 1)***. However, the processing of ILD has been shown to already occur at the level of the brainstem (for a review see ***Grothe et al. (2010)***; ***Yin et al. (2019))***. In contrast, the cortex has been associated with higher order processing ***Rauschecker et al. (1995)***; ***Rauschecker (1997, 1998)***; ***Zatorre and Belin (2001)***; ***Wessinger et al. (2001)***.

We were also surprised that the locus of activity did not include auditory processing areas in the brainstem. Although the 3D-EPI sequence that was modified for ISSS imaging (see Materials and methods) improves signal detection in subcortical areas, in contrast to most fMRI studies, we collected the fMRI data sagitally in order to employ completely silent, non-selective volume excitation. Nevertheless, a typical phase encoding direction approximately along the anterior-posterior commissure line was used (AP). Therefore, typical susceptibility-induced distortions and smearing of the signal occurred in AP direction, particularly in the brainstem. However, due to the narrow-banded water-selective excitation used, more off-resonance-induced signal drop-outs may have occurred, in addition to typical intra-voxel dephasing. While geometric distortion correction can mitigate the observed distortions and signal smearing to some extent, it cannot recover signal that may have been lost in brainstem regions. We therefore cannot exclude the possibility that brainstem activity was also relevant for our task, but that it was not possible to resolve due to high signal noise and signal drop-out. Additional studies with simpler tasks and analyses should be performed to improve the sequence and adapt the preprocessing to consistently measure the activation in the brainstem.

## Conclusion

We investigated the impact of variations in auditory cues based on different BRIRs of the same physical room on the neural processing of spatial acoustics during a distance discrimination task. We used the same room that subjects were in and had them perform an explicit distance task to improve ecological validity of our results. Although acoustic differences between distances and BRIR auralizations were small, they influenced the behavior of subjects and modified the multivariate neural activity pattern. Different auralizations of the same room can be decoded from brain activity. Changes in the brain activity patterns for distance and auralization were correlated with resulting variations in DRR. These variations in DRR are represented in posterior non-primary auditory cortices responsible for higher-level cognitive processing such as identification of sound location. This provides further evidence that populations of neurons located in the posterior AC are sensitive to DRR. It is possible that these regions directly extract DRR from auditory signals, or they may be extracted elsewhere and only integrated and combined with other cues in these posterior auditory cortical regions. As the posterior AC and the association cortex are known to be involved in higher-level cognition, the observed correlation with DRR in these regions supports the relevance of DRR for extracting abstract information about the sound source ***Rauschecker et al. (1995)***; ***Rauschecker (1997)***; ***Rauschecker and Tian (2000)***; ***Rauschecker (1998)***; ***Ahveninen et al. (2006)***. We demonstrate the use of RSA in determining the relevant acoustic properties in a complex auditory task. We are unaware of an auditory fMRI study that has used RSA in this manner before. We also developed MRI-compatible recording techniques and modified a 3D-EPI sequence to have quiescent periods for auditory imaging that can be used to measure brain activity ever few hundred milliseconds with a high isotropic resolution. We believe this sequence and recording equipment can benefit the larger auditory imaging community.

## Materials and methods

### Subjects

Initially, we collected the data from 20 pilot subjects. For the actual data acquisition we recruited 43 subjects. Of these, 13 were excluded due to poor data quality, initial problems with the setup, or because subjects had to abort the experiment. In total, the data from 17 female and 13 male subjects with normal hearing were included in the analyses (27 right-handed, 3 left-handed, age range: 19 to 34 years). All persons gave their informed consent to participate in the study and were monetarily compensated for their time at 12 € per hour. The experimental protocols were conducted in accordance with the Declaration of Helsinki and was approved by the independent ethics committee of the University of Munich (21-0035). Participants were not recruited if they were pregnant or had any contraindication for entry into the MRI bore.

### Stimuli and setup

Stimuli were originally anechoic sounds consisting of semantically neutral words with a fixed duration of 550 ms that were designed specifically for acoustic experiments in the MRI scanner ***Quadflieg et al. (2008)***. Sounds were subsequently convolved with one of three sets of BRIRs at one of two fixed positions within the environment. Three sets of BRIRs were created by capturing the acoustics of the magnet room ***(Figure 4a)*** with varying acoustic complexity. For each auralization of the MRI room, BRIRs were generated at two distances (see ***Figure 1c)***. Both distances were rightly lateralized with respect to the listener, as externalization is reported to be reduced for frontal sources compared to lateral ones ***Hassager et al. (2016)***. The farthest sound was placed in the right corner of the room to maximize the distance between the source and the receiver.

**Figure 4.**
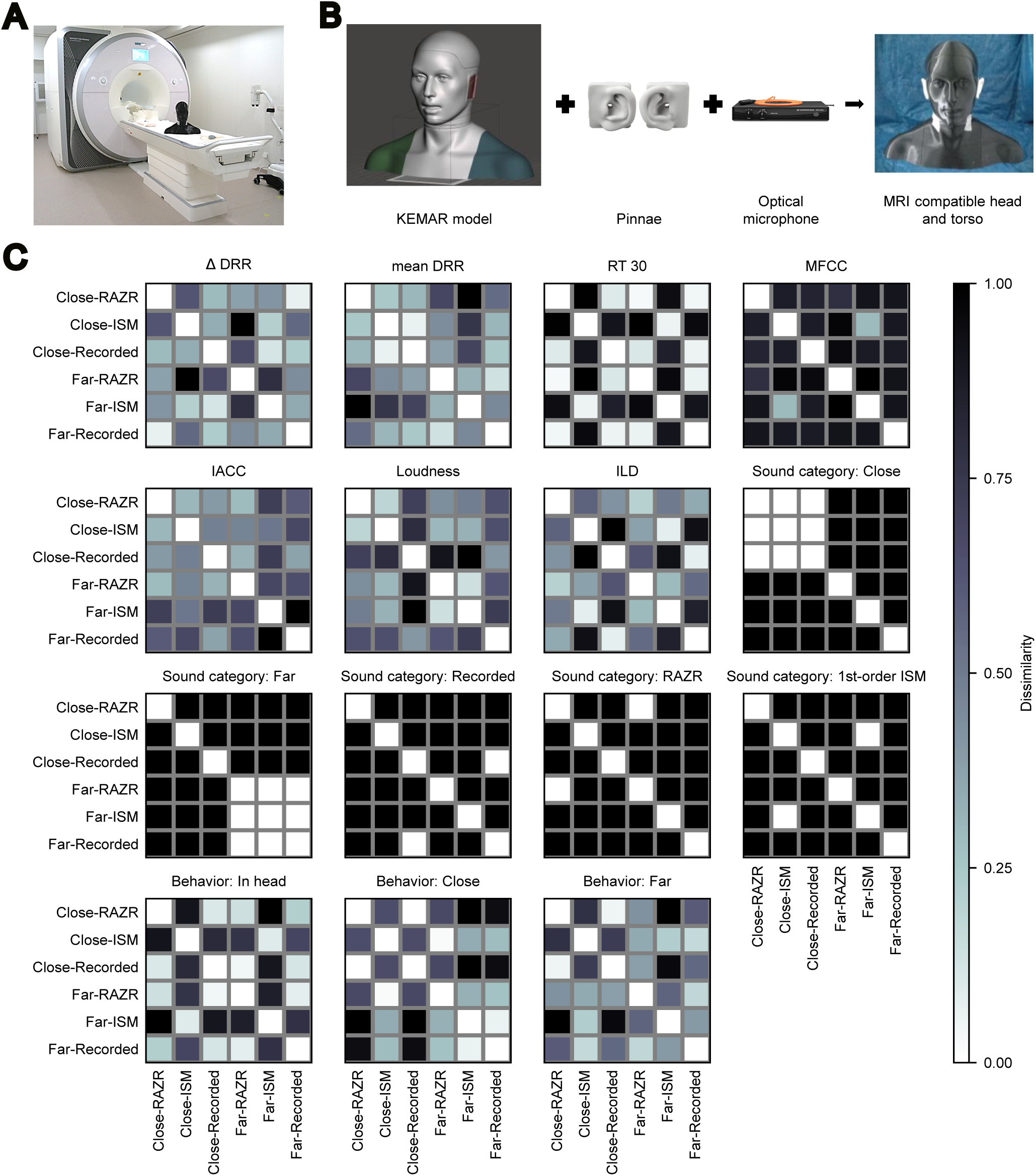
We conducted an RSA to examine the brain response to auralizations of the MRI scanner room that describe the acoustics with different degrees of accuracy. (A) The magnet room with the scanner and the MRI compatible head and torso simulator. (B) To record the magnet room we built an MRI compatible head and torso simulator that consisted of a KEMAR head and torso model ***Braren and Fels (2020)***, pinnae from a Brüel&Kjær dummy head and the microphone MO 2000 CU from Sennheiser. (Images from **Pin** (https://www.bksv.com/en/transducers/simulators/head-and-torso/pinnae-dz9770) and **Ford** (https://tyfordaudiovideo.blogspot.com/2013/06/sennheiser-mo-2000-industrial-optical.html)). (C) Model RDMs based on the acoustic parameters of the stimuli, the category of each stimulus and the behavioral response.

The first set of BRIRs was obtained by recording the acoustics of the magnet room. Since the magnetic fields of the scanner prohibit the usage of traditional dummy heads, we built an MRI compatible head and torso simulator. We combined a three-dimensional print of a KEMAR head and torso model by Braren and Fels ***Braren and Fels (2020)*** with pinnae from an existing Brüel&Kjær dummy head. Inside the simulator we positioned the microphone MO 2000 CU (Sennheiser, Germany) ***(Figure 4b)***. Headphones were connected to a Babyface Pro (RME, Germany) audio interface. Using the aforementioned MRI compatible head and torso, we recorded BRIRs inside the magnet room. The MRI cooling system and cold head were temporarily turned off during recording.

The second and third auralization were created using the offline version of the room acoustic simulator RAZR ***Wendt et al. (2014)***. The simulator combines an image source model for the specular reflections with a feedback delay network for modeling the diffuse late reverberation. The simulation (referred to as *RAZR*) was adjusted to match the acoustical properties of the recorded room by aligning RT30, geometry as well as source and receiver position while neglecting objects inside the room. For the third auralization (referred to as *1st-order ISM*), we simplified the above RAZR model from a third-order to a first-order image source model and disabled the feedback delay network.

This procedure resulted in six different BRIRs: two distances × three room auralizations. BRIRs were peak normalized. Additionally, stimuli were equalized to account for the headphone transfer function and in the case of the recorded BRIRs, the transfer function of the loudspeaker and microphone. All stimuli were pre-generated at a sampling rate of 44.1 kHz.

### Experimental procedure

The experiment started with a training phase with feedback, which was conducted in the same position as the final experiment. During training as well as the experiment, the task was to identify the perceived distance to the presented sound, classifying it as originating from the corner of the room (*Far*), at the level of their feet (*Close*), or very close to or inside their head (*In head*).

Stimuli were presented via MRI compatible electronic in-ear headphones (Model S14, Sennheiser, Germany). Words were presented in blocks of four, in which the same word and room simulation was maintained within each block. Subjects were correctly informed that the first sound of each block was located at the level of their feet (*Close*). For the remaining sounds the distance were randomly varied for a fixed set of eight randomizations that counterbalanced distance and room auralization. Subjects completed eight runs, with 9 trials for a total of 36 stimuli for each run. All runs were acquired in one session with short breaks between the runs. A large number of shorter runs was chosen to have a greater number of independent datasets for classification.

### fMRI acquisition

The experiment was performed on a Siemens Prisma 3 T MRI scanner using a 64-channel head coil (Siemens Healthineers, Erlangen, Germany). We implemented a custom multi-shot 3D-EPI sequence with ISSS imaging ***Schwarzbauer et al. (2006)***. ISSS is a sparse imaging method that allows for multiple image acquisitions after a silent period used for stimulus presentation. Within the silent period, the imaging gradients are not performed to remove the most dominant, high-pitched sequence noise. At the same time, the steady-state longitudinal magnetization is maintained by continued excitation at the same rate as in the loud period, thus avoiding T1-related signal changes that would otherwise occur.

The original ISSS method used a 2D single band EPI sequence. We extended it to a multi-shot 3D-EPI sequence with blipped-CAIPI parallel imaging ***Stirnberg and Stöcker (2021)*** and time-efficient water excitation ***Stirnberg et al. (2017)***. Thus, we achieve whole brain coverage at a high temporal resolution of 0.57 s volume repetition time (TR) along with 2.5 mm isotropic voxel size. With 3D-EPI, the silent period can be defined by any integer multiple of the TR per shot instead of the volume TR. Here, we used 88 × 51.81 ms. Furthermore, 3D-EPI allowed us to use spatially non-selective water excitation (no slice-selective gradient pulses) to substantially reduce non-pitched noise contributions. Finally, extremely quiet spoiler gradients were implemented in the silent periods by maximizing their duration between successive excitations (minimizing their amplitudes) while maintaining the same effective spoiling moments as in the loud periods. The final fMRI sequence parameters were: 225 mm×225 mm×150 mm field-of-view, sagittal slice orientation (phase encoding approximately along the anterior-posterior commissure line), ‘WE-rect 1-2-1’ water excitation***Norbeck et al. (2020)***, 16^◦^ flip angle, 1×6_*z*1_ blipped-CAIPI parallel imaging, 7/8 partial Fourier sampling (skipping only most decayed echoes > TE), 11 shots per volume including one initial “dummy shot” used for phase navigator acquisition, TE=30 ms.

The stimulus was presented within the silent period of 4.559 s, which was followed by 12 fMRI image volumes in a period of 6.84 s. In total, 542 image volumes were acquired per run, resulting in TA=8:43 min total acquisition time.

Between runs, two additional 3D-EPI images were acquired for subsequent geometric distortion correction, one with identical and one with inverted phase encoding, as well as a high-resolution T1-weighted anatomical MPRAGE image at 0.75 mm isotropic resolution (240 mm×240 mm×192 mm field-of-view, sagittal slice orientation, 12^◦^ flip angle, 2-fold GRAPPA parallel imaging, TE=2.17 ms, TR=2060 ms, TI=1040 ms, TA=6:35 min).

### Data analysis

#### Acoustic parameters

In order to quantify how the six auralizations (two distances in three room auralizations) differed from one another, we analyzed the acoustic parameters of the sounds presented. All acoustic parameters were computed using MATLAB 2021b ***MATLAB (2017)***. RT30 was determined based on the Schroeder integration method ***Schroeder (1965)***. To compute the IACC, we aligned the onset of the BRIRs based on the first peak reaching a magnitude of ten times the noise floor. Subsequently, the IACC was calculated on the aligned versions of the BRIRs as suggested by EN ISO 3382-1:2009 ***ISO Central Secretary (2009)***. For the determination of the DRR, the direct and reverberant part of the sound were extracted based on a time-window procedure. Direct sound and reverberation were separated by estimating the onset of the reverberation. Everything before this point is considered the direct sound, while the interval beyond is considered reverberant. The room acoustic simulator RAZR enables to obtain the direct and reverberant part, separately. Hence, we could compute the DRR for the simulation exactly. However, the recorded BRIRs require to estimate the separation between direct sound and the reverberant part based on the full BRIR. Hence, to compare the DRR among room auralizations we used the time-window procedure for all three room auralization. The exact separation in direct and reverberant part for the simulation was used to estimate the onset of the reverberation. The DRR value obtained via the time-window procedure was aligned with the value based on the exact separation from RAZR by varying the estimated onset of the reverberation. Finally, the point in time providing the closest match was used as the onset of the reverberation. Based on this alignment the duration of the direct sound was defined as 0.65 ms starting from the first peak of the BRIR that reached a magnitude of ten times the noise floor. To avoid artifacts, the filters were designed to start and end with with the decreasing and increasing part of a Hann-filter, respectively. Time to peak of the Hann-filter was 0.5 ms. The perceived loudness of the convolved sounds was computed according to ISO 532-2 (Moore-Glasberg) ***Moore and Glasberg (1996)***.

#### Behavioral data

The percentage of the different possible responses was examined for each room auralization. Correct responses were not calculated, as the additional possible *In head* response made an exact calculation of correct performance infeasible. Since responses were categorical, a multinomial logistic regression was used to model the response behavior. Logistic models predict the log-odds for the dependent variable based on a linear combination of the independent variables. The estimated coefficients describe how the independent variables influence the probability of the dependent variable to take a certain level, in our case *Close*, *Far* or *In head*. To test for a significant change in *In head* responses we compared the amount of *In head* responses with the sum of *Close* and *Far* responses, effectively reducing the regression to a binomial logistic regression with the dependent variables *In head* versus “not *In head*” i.e. *Close* or *Far*. The regression analysis was carried out using R ***R Core Team (2023)*** with the R package “mclogit” ***Elff (2022)***.

#### Preprocessing of imaging data

Results included in this manuscript come from preprocessing performed using fMRIPrep 23.1.3 ***(Esteban et al. (2019, 2018)*** RRID:SCR_016216), based on Nipype 1.8.6 ***(Gorgolewski et al. (2011)***; **Esteban et al.** (https://doi.org/10.5281/zenodo.5585697); RRID:SCR_002502).

##### Anatomical data preprocessing

The T1-weighted (T1w) image was corrected for intensity non-uniformity (INU) with N4BiasFieldCorrection ***(Tustison et al., 2010)***, distributed with ANTs (version unknown) ***(Avants et al., 2008***, RRID:SCR_004757), and used as T1w-reference throughout the workflow. The T1w-reference was then skull-stripped with a *Nipype* implementation of the antsBrainExtraction.sh workflow (from ANTs), using OASIS30ANTs as target template. Brain tissue segmentation of cerebrospinal fluid (CSF), white-matter (WM) and gray-matter (GM) was performed on the brain-extracted T1w using fast (FSL (version unknown), RRID:SCR_002823, ***Zhang et al., 2001)***. Brain surfaces were reconstructed using recon-all (FreeSurfer 7.3.2, RRID:SCR_001847, ***Dale et al., 1999)***, and the brain mask estimated previously was refined with a custom variation of the method to reconcile ANTs-derived and FreeSurfer-derived segmentations of the cortical gray-matter of Mindboggle (RRID:SCR_002438, ***Klein et al., 2017a)***. Volume-based spatial normalization to one standard space (MNI152NLin2009cAsym) was performed through nonlinear registration with antsRegistration (ANTs (version unknown)), using brain-extracted versions of both T1w reference and the T1w template. The following template was selected for spatial normalization and accessed with *TemplateFlow* (23.0.0, ***Ciric et al., 2022)***: *ICBM 152 Nonlinear Asymmetrical template version 2009c* [***Fonov et al. (2009)***, RRID:SCR_008796; TemplateFlow ID: MNI152NLin2009cAsym].

##### Functional data preprocessing

For each of the 8 BOLD runs found per subject (across all tasks and sessions), the following preprocessing was performed. First, a reference volume and its skull-stripped version were generated using a custom methodology of *fMRIPrep*. Head-motion parameters with respect to the BOLD reference (transformation matrices, and six corresponding rotation and translation parameters) are estimated before any spatiotemporal filtering using mcflirt (FSL, ***Jenkinson et al., 2002)***. The BOLD time-series (including slice-timing correction when applied) were resampled onto their original, native space by applying the transforms to correct for head-motion. These resampled BOLD time-series will be referred to as *preprocessed BOLD in original space*, or just *preprocessed BOLD*. The BOLD reference was then co-registered to the T1w reference using bbregister (FreeSurfer) which implements boundary-based registration ***(Greve and Fischl, 2009)***. Co-registration was configured with six degrees of freedom. Several confounding time-series were calculated based on the *preprocessed BOLD*: framewise displacement (FD), DVARS and three region-wise global signals. FD was computed using two formulations following Power (absolute sum of relative motions, ***Power et al. (2014))*** and Jenkinson (relative root mean square displacement between affines, ***Jenkinson et al. (2002))***. FD and DVARS are calculated for each functional run, both using their implementations in *Nipype* (following the definitions by ***Power et al., 2014)***. The three global signals are extracted within the CSF, the WM, and the whole-brain masks. Additionally, a set of physiological regressors were extracted to allow for component-based noise correction (*CompCor*, ***Behzadi et al., 2007)***. Principal components are estimated after high-pass filtering the *preprocessed BOLD* time-series (using a discrete cosine filter with 128s cut-off) for the two *CompCor* variants: temporal (tCompCor) and anatomical (aCompCor). tCompCor components are then calculated from the top 2% variable voxels within the brain mask. For aCompCor, three probabilistic masks (CSF, WM and combined CSF+WM) are generated in anatomical space. The implementation differs from that of Behzadi et al. in that instead of eroding the masks by 2 pixels on BOLD space, a mask of pixels that likely contain a volume fraction of GM is subtracted from the aCompCor masks. This mask is obtained by dilating a GM mask extracted from the FreeSurfer’s *aseg* segmentation, and it ensures components are not extracted from voxels containing a minimal fraction of GM. Finally, these masks are resampled into BOLD space and binarized by thresholding at 0.99 (as in the original implementation). Components are also calculated separately within the WM and CSF masks. For each CompCor decomposition, the *k* components with the largest singular values are retained, such that the retained components’ time series are sufficient to explain 50 percent of variance across the nuisance mask (CSF, WM, combined, or temporal). The remaining components are dropped from consideration. The head-motion estimates calculated in the correction step were also placed within the corresponding confounds file. The confound time series derived from head motion estimates and global signals were expanded with the inclusion of temporal derivatives and quadratic terms for each ***(Satterthwaite et al., 2013)***. Frames that exceeded a threshold of 0.5 mm FD or 1.5 standardized DVARS were annotated as motion outliers. Additional nuisance timeseries are calculated by means of principal components analysis of the signal found within a thin band (*crown*) of voxels around the edge of the brain, as proposed by ***(Patriat et al., 2017)***. The BOLD time-series were resampled into standard space, generating a *preprocessed BOLD run in MNI152NLin2009cAsym space*. First, a reference volume and its skull-stripped version were generated using a custom methodology of *fMRIPrep*. All resamplings can be performed with *a single interpolation step* by composing all the pertinent transformations (i.e. head-motion transform matrices, susceptibility distortion correction when available, and co-registrations to anatomical and output spaces). Gridded (volumetric) resamplings were performed using antsApplyTransforms (ANTs), configured with Lanczos interpolation to minimize the smoothing effects of other kernels ***(Lanczos, 1964)***. Non-gridded (surface) resamplings were performed using mri_vol2surf (FreeSurfer).

Many internal operations of *fMRIPrep* use *Nilearn* 0.10.1 ***(Abraham et al., 2014***, RRID:SCR_001362), mostly within the functional processing workflow. For more details of the pipeline, see the section corresponding to workflows in *fMRIPrep*’s documentation.

Subsequently, the images were smoothed with a kernel of 8 mm full-width at half maximum (FWHM) using the SPM12 toolbox from MATLAB 2021b ***MATLAB (2017)***.

#### Univariate analysis

A univariate analysis was conducted using the SPM12 (7771) toolbox with MATLAB 2021b ***MATLAB (2017)***. A GLM was created with regressors for each of the six BRIRs. These were modeled as events time-locked to the sound onset. The Canonical Hemodynamic Response Function was used to model the hemodynamic effects. The regressors of interest were first created as if the fMRI data were collected with continuous sampling and then modified to reflect the silent periods without image data collection. Additionally, six motion regressors, three for translation and three for rotation, modeled the head-movement related changes in the MRI signal. A within-subject ANOVA with multiple levels was carried out to assess the main effects at the group level.

#### Decoding

Three linear support vector classifiers with L1-penalty were trained on the activity maps obtained from the univariate analysis to decode each room auralization from brain activity data. We applied the decoding scheme ‘FREMclassifier’ of the python package nilearn ***Abraham et al. (2014)***. As the noise within one run is autocorrelated, the activity maps were grouped according to the runs for each subject. Crossvalidation was performed, and the mean classification accuracy was calculated for each subject across folds. After testing for normality via a Lilliefors test (*Recorded*: *D* = 0.09, *p* = 0.72; *RAZR*: *D* = 0.11, *p* = 0.53; *1st-order ISM*: *D* = 0.11, *p* = 0.49;), we conducted a t-test to assess whether the average of these mean classification accuracies was significantly larger than chance.

Chance accuracy was defined as the average across chance levels per subject. A permutation test with *N* = 1000 was run for each subject to determine the chance accuracy at the subject level.

The searchlight analysis was based on three one-versus-rest support vector machines with L1-penalty. The support vector machines were combined in a one-versus-rest strategy for multiclass classification. The analysis was performed using the software package nilearn from python ***Abraham et al. (2014)***. The searchlight radius was set to 5.6 mm and the performance was evaluated based on the classification accuracy. To determine significance against chance level at the group level, we conducted a permutation test with 5000 permutations utilizing the MATLAB toolbox SnPM ***SnP (2004)***. As the occurrence of each room auralization was equalized across runs, the chance level of the combined classifiers was 0.33. A pseudo *t* statistics was used to reduce noise in the statistical maps. This pseudo *t* involves a local pooling of the variance i.e. averaging the variance estimates in the local neighborhood of the voxels (for details see ***SnP (2004))***.

#### Representational similarity analysis

The representational similarity analysis was conducted with the python package rsatoolbox ***Lu and Ku (2020)***. RSA relates neuroscientific models and/or behavior to brain activity. For more details refer to ***Kriegeskorte et al. (2008)***. In a first step, we conducted a GLM on the subject level to determine the activity map for each combination of room and distance. Subsequently, we computed the crossvalidated squared Mahalanobis distance for each pair of activity maps ***Diedrichsen and Kriegeskorte (2017)*** resulting in a dissimilarity matrix (RDM, as in ***Figure 4c)*** for each subject. The matrices were averaged over all subjects to obtain one RDM characterizing the measured activity pattern.

To identify how acoustic cues relate to brain activity, the acoustic parameters that are relevant for ADP ***Kolarik et al. (2016)***; ***Zahorik et al. (2005)*** were calculated (see Materials and methods, **subsubsection** Acoustic parameters). For each parameter, dissimilarity matrices were created.

The matrices based on DRR, RT30, MFCC and IACC were generated by directly computing the respective parameter for each BRIR per condition. In contrast, perceived loudness and ILD were calculated for each of the convolved sounds and these values were then averaged across sounds per condition.

The parameters RT30, IACC and perceived loudness were computed for the frequencies 250 Hz, 500 Hz, 750 Hz, 1 kHz, 1.5 kHz and 2 kHz. Hence, for these parameters we obtained a vector with six values for each of the six conditions. The dissimilarity metric between conditions based on these acoustic cues was defined as one minus the Pearson correlation of the respective frequency vectors.

For ILD, we obtained one value per condition. The dissimilarity measure was based on the Euclidean distance between the ILD values of two conditions. Regarding MFCC, dissimilarity was defined as one minus the Pearson correlation of the mel-coefficients for each BRIR. For DRR, we defined two dissimilarity metrics. First, DRR was computed for each channel resulting in two values per condition. Second, we calculated either the average DRR of the two channels (*mean DRR*) or the difference between the right and the left channel (Δ*DRR*). Finally, for both metrics we determined the dissimilarity between conditions by computing the euclidean distance between the values of each condition.

For the behavioral models, the percentage of *In head*, *Close* and *Far* responses was computed per condition. Dissimilarity of two conditions was determined by computing the Euclidean distance between the percentage values. Thus, we obtain three behavioral RDMs, one for each response type.

The categorical models were based on the stimulus category. We assigned a low dissimilarity of 0 if the stimuli of the two conditions belonged both to a specific distance or auralization type and 1 otherwise. Thus, we obtain five additional model RDMs, two based on distance, namely *Close* and *Far* and three based on room auralization referred to as *In head*, *RAZR* and *1st-order ISM*.

The resulting model RDMs were normalized between 0 and 1 so that the weights reflected relative contributions and not an inverse of the absolute values of the parameters themselves. ***Figure 4c*** depicts the model RDMs for each acoustic parameter.

To investigate how acoustic parameters may be related to the brain activity we computed the whitened-RDM Pearson correlation between each model RDM and the RDM based on the measured brain activity. For details regarding the correlation parameter, see ***Diedrichsen and Kriegeskorte (2017)***. The weighted models predict the data RDM as a linear weighted sum of the individual model RDMs. An optimization algorithm based on the Broyden–Fletcher–Goldfarb–Shanno procedure was applied to fit the weighted models to the data. Fit accuracy was assessed using crossvalidation. To obtain an estimate of the variance, we conducted bootstrapping with 1000 samples.

## Acknowledgments

We thank Stefan Fichna, Thomas Eggert and Thomas Stephan for their comments on the analysis, Hans Hintermaier for his support with the MRI compatible head and torso and Siegfried Gündert for his assistance and the provided software concerning the measurement of the IRs.

## Data availability

Data and Code will be made available upon acceptance on G-node (https://gin.g-node.org/) and gitlab (https://gitlab.lrz.de/).

## Funding

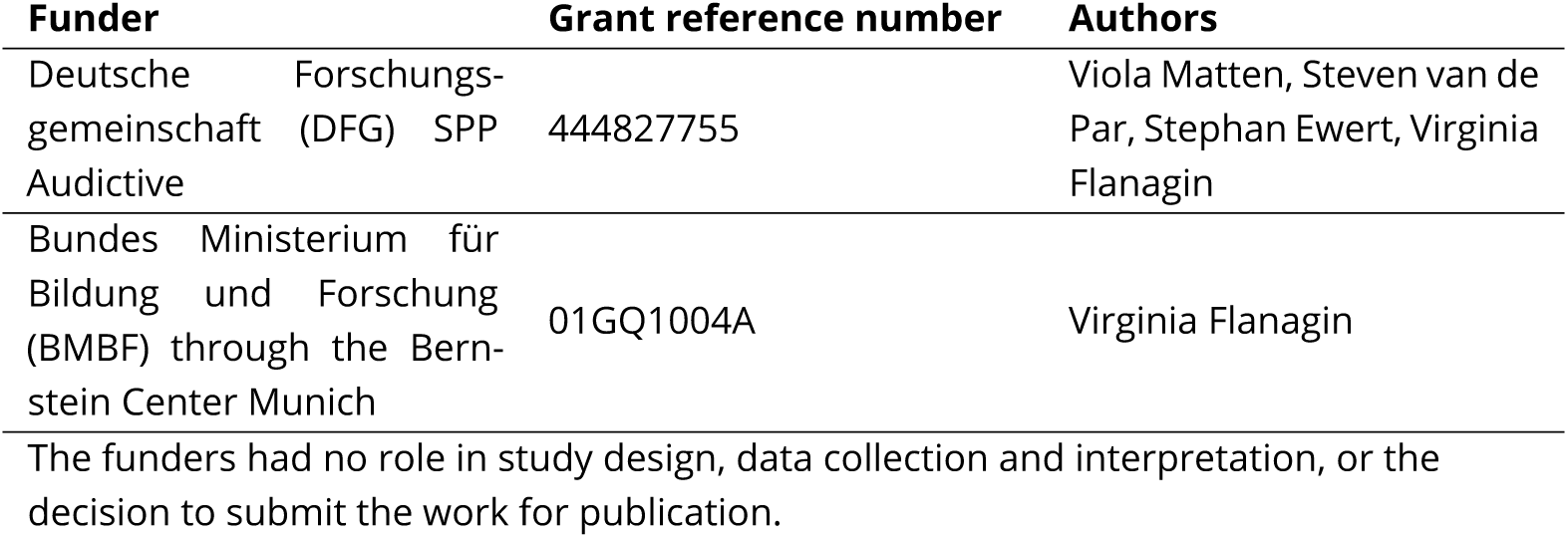

## Author contributions

Viola G Matten, Conceptualization, Methodology, Software, Formal analysis, Investigation, Resources, Writing – original draft preparation, Writing – review & editing; Visualisation; Rüdiger Stirnberg, Software, Methodology, Writing – review & editing; Steven van de Par, Stephan D Ewert, Conceptualization, Methodology, Software, Validation, Resources, Writing – original draft preparation; Writing – review & editing, Funding acquisition; Virginia L Flanagin, Conceptualization, Methodology, Software, Resources, Writing – review & editing; Visualisation, Supervision, Project administration, Funding acquisition;

## Ethics

The experimental protocols were conducted in accordance with the Declaration of Helsinki and were approved by the independent ethics committee of the University of Munich (21-0035). Participants were not recruited if they were pregnant or had any contraindication for entry into the MRI bore.

## Appendix 1

### Acoustic analysis of stimuli

This supplement provides more detailed information on how the acoustic analysis of the stimuli used in this study was performed. In addition, the analysis will show how acoustic cues vary across the different distances (*Close* and*Far*), as well as across the three different auralizations that were used (*Recorded, RAZR* and *1st-order ISM*). Modifying the complexity of the room auralization changed the acoustic cues in the studied dimensions distance and room auralization.

### Reverberation time, RT30

The reverberation time, RT30, of the recorded room ranges between 0.4 s and 0.45 s for audible frequencies and matches the RT30 of a typical office or residential building ***Reinhart et al. (2016)***. While RT30 was in a similar range for the recorded BRIRs compared to the full *RAZR* simulation, we observed a decrease for the *1st-order ISM*. This decrease can be attributed to the missing late reverberation in the *1st-order ISM*. Between the distances RT30 was found to be highly similar.

### Direct-to-reverberant ratio, DRR

In general, the direct-to-reverberant ratio, DRR is larger for the right channel than the left channel as sound sources were simulated on the right side of the subject. As expected DRR is smaller for sounds located further away from the listener. However, with an average of 2.8 ±2.2 dB DRR differences between distances are below JNDs ***Zahorik (2002b)***; ***Larsen et al. (2008)***. Between auralizations differences in DRR are larger with an average of 4.5 ± 2.1 dB and a maximum differences of 9 dB between the *1st-order ISM* and *RAZR*. This increase in DRR for the *1st-order ISM* is expected due to omitting parts of the reverberant tail.

Additionally, DRR of the recorded BRIRs is slightly larger than for the full RAZR simulation. We designed the full RAZR simulation such that it models the acoustics of the recorded room as accurate as possible but neglects objects inside the room. We hypothesize that the MRI scanner introduces additional very early reflections and thereby causes the observed change in DRR.

Note also that the subject was positioned such that there was a direct acoustic path from the source to the right ear while the left ear was in the acoustic shadow of the head. For that reason the direct sound component was more pronounced in the right ear leading to generally higher DRR values.

### Interaural level differences, ILDs

***Appendix 2 — Figure 1*** visualizes the average interaural level difference, ILD, for all sounds presented per condition. We observed an increase in ILDs as the complexity of the auralization increased. Surprisingly, we found a negative ILD for the recorded BRIRs although sounds are presented on the right side.

Plotting the level of the the early reflections and the reverberation separately from the level of the direct sound, suggests that this observation was caused by early reflections and the reverberation. We find that the level of the direct sound was larger for the right channel than the left channel, as expected ***(Appendix 2 — Figure 1b)***. The same applies to the reverberation level for the two simulations ***(Appendix 2 — Figure 1c)***. However, in the case of the recorded BRIRs the level of reverberation is larger for the left than for the right channel. We hypothesize that this effect is caused by reflections and diffraction at the scanner bore which was not included in the room acoustic simulations.

### Mel frequency cepstral coefficients, MFCC

The MFCCs are a perceptually relevant representation of the spectral shape of an acoustic stimulus, that is closely related to the timbre of a sound. The successive components represent the spectral shape on a MEL-frequency scale in terms of Discrete Cosine Transforms. The first coefficient represents overall level, the second, the strength of mid-frequency reduction of spectral energy. The coefficients provide an orthogonal decomposition of the MEL-frequency spectrum, with ever increasing detail with increasing coefficient number. MFCCs have been used extensively to represent the acoustic spectrum in a compact manner to allow pattern classifiers to recognize speakers identity or to recognize spoken words. Between the conditions we observed similar values for the first coefficient but considerable differences for the higher order coefficient ***(Appendix 2 — Figure 1d)***. Hence, the data suggests that variations in level are small while the spectral composition differs between the conditions.

### Interaural cross correlation, IACC

The IACC is a parameter that measures the similarity of the acoustic waveform arriving at both ears. Specifically, in reverberant conditions, the IACC will drop in value due to incoherent reflections that arrive from different directions. It has been shown that the IACC correlates to perceived source width. In the evaluation of concert-halls this parameter predicts Apparent Source Width and allows predicting preferences of concert halls ***Hidaka et al. (1995)***. The IACC is depicted in ***Appendix 2 — Figure 1e***. The two auralizations including direct sound together with early and late reverberation, *Recorded* and *RAZR*, provide perceptually similar IACCs ***Culling et al. (2001)***. For the *1st-order ISM* the IACC is increased compared to *Recorded* and *RAZR*. The observation is consistent with the fact that we include fewer reflections in the *1st-order ISM*.

Furthermore, when distance increases, the direct sound will reduce in level, while the reverberant sound field component will stay more or less at the same level. Thus, it is expected that with increasing distance, IACC will drop. This can be seen for *RAZR* at frequencies of 1 kHz and beyond. However, for low frequencies we observe the opposite behavior. While for *RAZR* the difference between *Close* and *Far* is close to JNDs for IACC ***Culling et al. (2001)***, the effect is more pronounced for the *1st-order ISM*. The increase in IACC for the condition *Far* might result from the proximity of the far sound source to the corner of the room. The placement close to the wall can increase the coherence for the direct sound and the first reflection. The presence of late reverberation for *RAZR*, in contrast to the *1st-order ISM*, might explain why the effect is stronger for the *1st-order ISM*. For the recorded BRIRs the presence of the scanner bore might introduce additional reflections that further reduce the IACC. These additional reflections differ most likely between *Close* and *Far*, and might explain why for *Recorded* the IACC is smaller for the condition *Far* than the condition *Close*.

### Perceived loudness

With reduced auralization accuracy, we observed a slight decrease in average loudness ***(Appendix 2 — Figure 1f)***. As expected, loudness was elevated for the condition *Close* compared to *Far*. However, in both dimensions, distance and room auralization, differences were below JNDs ***You and Jeon (2008)***.

## Appendix 2

**Appendix 2—figure 1.**
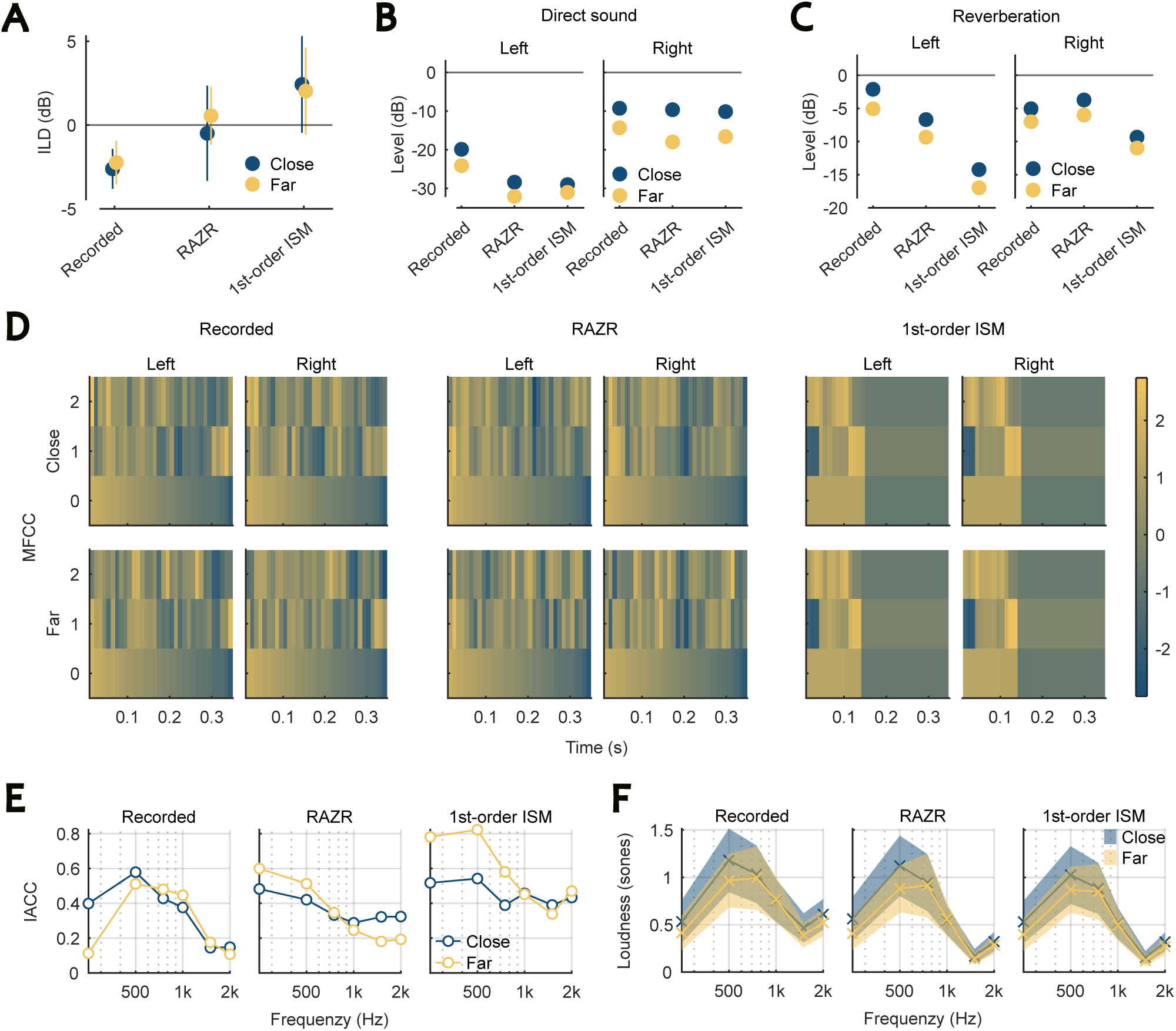
Acoustic analysis of the stimuli in the two dimensions auralization and distance. Stimuli differ mainly between auralizations and less between distances. (A) The ILD averaged over sounds per condition. Errorbars display standard deviation from the mean. (B-D) The level of the direct sound and the reverberation as well as the MFCC were calculated for the left and the right channel of each BRIR per condition. (E) The IACC between the left and the right channel of the BRIRs was determined for each condition. (F) The predicted loudness averaged over sounds per condition. Errors represent standard deviation from the mean.

## Appendix 3

**Appendix 3—table 1.** Significant clusters obtained in the univariate analysis. The statistical map has been thresholded at *p* = 0.05 FWE. Only cluster with more than 100 voxels were considered significant. Coordinates are given in MNI152 space.

| Cluster ID | X (mm) | Y (mm) | Z (mm) | Peak $t$ statistic | Volume (mm<sup>3</sup>) |
| --- | --- | --- | --- | --- | --- |
| 1 | -56.5 | -17.5 | 4.0 | 21.5 | 38687 |
| 2 | 61.0 | -7.5 | -1.0 | 20.1 | 38859 |
| 3 | -6.5 | 5.0 | 54.0 | 12.5 | 17906 |
| 4 | 31.0 | 27.5 | -1.0 | 10.9 | 4453 |
| 5 | -31.5 | 25.0 | 4.0 | 10.6 | 3390 |
| 6 | -39.0 | -22.5 | 54.0 | 10.1 | 17218 |
| 7 | 48.5 | 2.5 | 49.0 | 9.4 | 3968 |
| 8 | 43.5 | 15.0 | 26.5 | 8.6 | 4250 |
| 9 | 6.0 | -20.0 | -1.0 | 8.1 | 6125 |

## Appendix 4

**Appendix 4—table 1.** Data of the significant clusters obtained with the searchlight MVPA. The statistical map has been thresholded at *p* = 0.05 FWE and only clusters with more than 50 voxels were considered significant. Coordinates are given in MNI152 space.

| Cluster ID | X (mm) | Y (mm) | Z (mm) | peak pseudo-t | Volume (mm<sup>3</sup>) |
| --- | --- | --- | --- | --- | --- |
| 1 | -64.0 | -43.5 | 14.0 | 6.8 | 1750 |

## Appendix 5

**Appendix 5—table 1.** Prediction accuracies and statistics of all models studied with the whole brain RSA. Three distinct model types were investigated that characterized the conditions based on the acoustic parameters, the stimulus category i.e. whether sounds of that condition belonged to the same room auralization or distance and the average behavioral response in each condition. The models were correlated with the activity of the entire brain. Errors were obtained using bootstrapping (*N* = 1000). One-sided, uncorrected *p* values were computed testing whether the model accuracies differed significantly from zero and the noise ceiling, respectively.

| Model type | Model name | Accuracy | $p$ value (against 0) | $p$ value (against noise ceiling) |
| --- | --- | --- | --- | --- |
| Weighted models | DRR only | $0.18 \pm 0.07$ | 0.02 | 0.95 |
| | all acoustic parameters | $0.16 \pm 0.08$ | 0.05 | 0.91 |
| Acoustic parameters | mean DRR | $0.14 \pm 0.11$ | 0.14 | 0.76 |
| | $\Delta$ DRR | $0.07 \pm 0.15$ | 0.33 | 0.57 |
| | RT30 | $0.04 \pm 0.10$ | 0.374 | 0.286 |
| | MFCC | $0.03 \pm 0.13$ | 0.40 | 0.35 |
| | IACC | $-0.08 \pm 0.15$ | 0.68 | 0.21 |
| | Loudness | $0.003 \pm 0.13$ | 0.49 | 0.28 |
| | ILD | $-0.026 \pm 0.115$ | 0.587 | 0.20 |
| Category | Recorded | $-0.06 \pm 0.07$ | 0.78 | 0.08 |
| | RAZR | $0.02 \pm 0.06$ | 0.38 | 0.13 |
| | 1st-order ISM | $0.01 \pm 0.05$ | 0.39 | 0.13 |
| | Close | $0.09 \pm 0.1$ | 0.19 | 0.46 |
| | Far | $0.13 \pm 0.09$ | 0.12 | 0.65 |
| Behavior | In head | $0.04 \pm 0.12$ | 0.38 | 0.37 |
| | Close | $0.09 \pm 0.14$ | 0.28 | 0.59 |
| | Far | $0.10 \pm 0.12$ | 0.22 | 0.58 |

## Appendix 6

**Appendix 6—table 1.** Data of the significant clusters obtained with the searchlight RSA. The statistical map has been thresholded at *p* = 0.05 FWE and only clusters with more than 50 voxels were considered significant. Coordinates are given in MNI152 space.

| Cluster ID | X (mm) | Y (mm) | Z (mm) | Peak $t$ statistic | Volume (mm<sup>3</sup>) |
| --- | --- | --- | --- | --- | --- |
| 1 | -69.0 | -15.0 | -13.5 | 7.1 | 3453 |
| 2 | -69.0 | -45.0 | 11.5 | 7.0 | 5312 |
| 3 | 61.0 | -12.5 | -13.5 | 6.2 | 2546 |

## References

Chapter 46 - Nonparametric Permutation Tests for Functional Neuroimaging. In: Frackowiak RSJ, Friston KJ, Frith CD, Dolan RJ, Price CJ, Zeki S, Ashburner JT, Penny WD, editors. Human Brain Function (Second Edition), second ed. Burlington: Academic Press; 2004.p. 887–910. https://www.sciencedirect.com/science/article/pii/B9780122648410500482, doi: 10.1016/B978-012264841-0/50048-2.

Symmetrical Pinnea DZ-9770 + DZ-9769 from Brüel and Kjær; https://www.bksv.com/en/transducers/simulators/head-and-torso/pinnae-dz9770. Accessed 07 Mai 2024.

Abraham A, Pedregosa F, Eickenberg M, Gervais P, Mueller A, Kossaifi J, Gramfort A, Thirion B, Varoquaux G. Machine learning for neuroimaging with scikit-learn. Frontiers in Neuroinformatics. 2014; 8. https://www.frontiersin.org/articles/10.3389/fninf.2014.00014/full, doi: 10.3389/fninf.2014.00014.

Ahveninen J, Jääskeläinen IP, Raij T, Bonmassar G, Devore S, Hämäläinen M, Levänen S, Lin FH, Sams M, Shinn-Cunningham BG, et al. Task-modulated “what” and “where” pathways in human auditory cortex. Proceedings of the National Academy of Sciences. 2006; 103(39):14608–14613.

Ahveninen J, Kopčo N, Jääskeläinen IP. Psychophysics and neuronal bases of sound localization in humans. Hearing Research. 2014; 307:86–97. https://www.sciencedirect.com/science/article/pii/S0378595513001792, doi: 10.1016/j.heares.2013.07.008.

Arend JM, Ramírez M, Liesefeld HR, Pörschmann C. Do near-field cues enhance the plausibility of non-individual binaural rendering in a dynamic multimodal virtual acoustic scene? Acta Acustica. 2021; 5:55.

Avants BB, Epstein CL, Grossman M, Gee JC. Symmetric diffeomorphic image registration with cross-correlation: Evaluating automated labeling of elderly and neurodegenerative brain. Medical Image Analysis. 2008; 12(1):26–41. http://www.sciencedirect.com/science/article/pii/S1361841507000606, doi: 10.1016/j.media.2007.06.004.

Begault DR, Wenzel EM, Anderson MR. Direct comparison of the impact of head tracking, reverberation, and individualized head-related transfer functions on the spatial perception of a virtual speech source. Journal of the audio Engineering Society. 2001; 49(10):904–916.

Behzadi Y, Restom K, Liau J, Liu TT. A component based noise correction method (CompCor) for BOLD and perfusion based fMRI. NeuroImage. 2007; 37(1):90–101. http://www.sciencedirect.com/science/article/pii/S1053811907003837, doi: 10.1016/j.neuroimage.2007.04.042.

Best V, Baumgartner R, Lavandier M, Majdak P, Kopčo N. Sound Externalization: A Review of Recent Research. Trends in hearing. 2020 09; 24. doi: 10.1177/2331216520948390.

Brandenburg K, Klein F, Neidhardt A, Sloma U, Werner S. In: Blauert J, Braasch J, editors. Creating Auditory Illusions with Binaural Technology Cham: Springer International Publishing; 2020. p. 623–663. 10.1007/978-3-030-00386-9_21, doi: 10.1007/978-3-030-00386-9_21.

Braren HS, Fels J, A High-Resolution Head-Related Transfer Function Data Set and 3D-Scan of KEMAR; 2020. https://publications.rwth-aachen.de/record/807373, doi: 10.18154/RWTH-2020-11307.

Brechmann A, Baumgart F, Scheich H. Sound-level-dependent representation of frequency modulations in human auditory cortex: a low-noise fMRI study. Journal of Neurophysiology. 2002; 87(1):423–433.

Bregman AS. Auditory Scene Analysis: The Perceptual Organization of Sound. The MIT Press; 1990. 10.7551/mitpress/1486.001.0001, doi: 10.7551/mitpress/1486.001.0001.

Bronkhorst AW, Houtgast T. Auditory distance perception in rooms. Nature. 1999; 397(6719):517–520.

Bronkhorst A. Effect of stimulus properties on auditory distance perception in rooms. Physiological and Psychological Bases of Auditory Function. 2001; p. 184–191.

Brunetti M, Belardinelli P, Caulo M, DelGratta C, Della Penna S, Ferretti A, Lucci G, Moretti A, Pizzella V, Tartaro A, Torquati K, Belardinelli M, Romani GL. Human brain activation during passive listening to sounds from different locations: An fMRI and MEG study. Human brain mapping. 2005 12; 26:251–61. doi: 10.1002/hbm.20164.

Brungart DS, Durlach NI, Rabinowitz WM. Auditory localization of nearby sources. II. Localization of a broadband source. The Journal of the Acoustical Society of America. 1999 10; 106(4):1956–1968. 10.1121/1.427943, doi: 10.1121/1.427943.

Butler RA, Levy ET, Neff WD. Apparent distance of sounds recorded in echoic and anechoic chambers. Journal of Experimental Psychology: Human Perception and Performance. 1980; 6(4):745.

Catic J, Santurette S, Buchholz JM, Gran F, Dau T. The effect of interaural-level-difference fluctuations on the externalization of sound. The Journal of the Acoustical Society of America. 2013; 134:1232–1241. doi: 10.1121/1.4812264.

Catic J, Santurette S, Dau T. The role of reverberation-related binaural cues in the externalization of speech. The Journal of the Acoustical Society of America. 2015; 138(2):1154–1167.

Ciric R, Thompson WH, Lorenz R, Goncalves M, MacNicol E, Markiewicz CJ, Halchenko YO, Ghosh SS, Gorgolewski KJ, Poldrack RA, Esteban O. TemplateFlow: FAIR-sharing of multi-scale, multi-species brain models. Nature Methods. 2022; 19:1568–1571. doi: 10.1038/s41592-022-01681-2.

Coleman PD. An analysis of cues to auditory depth perception in free space. Psychological bulletin. 1963; 60(3):302.

Culling JF, Colburn HS, Spurchise M. Interaural correlation sensitivity. The Journal of the Acoustical Society of America. 2001; 110(2):1020–1029.

Dale AM, Fischl B, Sereno MI. Cortical Surface-Based Analysis: I. Segmentation and Surface Reconstruction. NeuroImage. 1999; 9(2):179–194. http://www.sciencedirect.com/science/article/pii/S1053811998903950, doi: 10.1006/nimg.1998.0395.

Demirkaplan Ö, Hacıhabiboğlu H. Effects of interpersonal familiarity on the auditory distance perception of level-equalized reverberant speech. Acta Acustica. 2020; 4(6):26.

Deouell LY, Heller AS, Malach R, D’Esposito M, Knight RT. Cerebral Responses to Change in Spatial Location of Unattended Sounds. Neuron. 2007; 55(6):985–996. https://www.sciencedirect.com/science/article/pii/S0896627307006484, doi: 10.1016/j.neuron.2007.08.019.

Dewey RS, Hall DA, Plack CJ, Francis ST. Comparison of continuous sampling with active noise cancelation and sparse sampling for cortical and subcortical auditory functional MRI. Magnetic resonance in medicine. 2021; 86(5):2577–2588.

Diedrichsen J, Kriegeskorte N. Representational models: A common framework for understanding encoding, pattern-component, and representational-similarity analysis. PLoS computational biology. 2017; 13(4).

Duda RO, Martens WL. Range dependence of the response of a spherical head model. The Journal of the Acoustical Society of America. 1998 11; 104(5):3048–3058. 10.1121/1.423886, doi: 10.1121/1.423886.

Elff M. mclogit: Multinomial Logit Models, with or without Random Effects or Overdispersion; 2022, https://CRAN.R-project.org/package=mclogit, r package version 0.9.6.

Esteban O, Blair R, Markiewicz CJ, Berleant SL, Moodie C, Ma F, Isik AI, Erramuzpe A, Kent M James D and- Goncalves, DuPre E, Sitek KR, Gomez DEP, Lurie DJ, Ye Z, Poldrack RA, Gorgolewski KJ. fMRIPrep. Software. 2018; doi: 10.5281/zenodo.852659.

Esteban O, Markiewicz C, Blair RW, Moodie C, Isik AI, Erramuzpe Aliaga A, Kent J, Goncalves M, DuPre E, Snyder M, Oya H, Ghosh S, Wright J, Durnez J, Poldrack R, Gorgolewski KJ. fMRIPrep: a robust preprocessing pipeline for functional MRI. Nature Methods. 2019; 16:111–116. doi: 10.1038/s41592-018-0235-4.

Esteban O, Markiewicz CJ, Burns C, Goncalves M, Jarecka D, et al. nipy/nipype: 1.7.0. Zenodo. https://doiorg/105281/zenodo5585697; Deposited 20 October 2021.

Fonov V, Evans A, McKinstry R, Almli C, Collins D. Unbiased nonlinear average age-appropriate brain templates from birth to adulthood. NeuroImage. 2009; 47, Supplement 1:S102. doi: 10.1016/S1053-8119(09)70884-5.

Ford T, Sennheiser MO 2000 Industrial Optical Analog Microphone; https://tyfordaudiovideo.blogspot.com/2013/06/sennmo-2000-industrial-optical.html. Accessed 27 September 2024.

Foster JJ, Ling S. Feature-Based Attention Multiplicatively Scales the fMRI-BOLD Contrast-Response Function. Journal of Neuroscience. 2022; 42(36):6894–6906. https://www.jneurosci.org/content/42/36/6894, doi: 10.1523/JNEUROSCI.0513-22.2022.

Gamble EA. Minor studies from the psychological laboratory of Wellesley College: Intensity as a criterion in estimating the distance of sounds. Psychological Review. 1909; 16(6):416.

Gandhi SP, Heeger DJ, Boynton GM. Spatial attention affects brain activity in human primary visual cortex. Proceedings of the National Academy of Sciences. 1999; 96(6):3314–3319.

Gorgolewski K, Burns CD, Madison C, Clark D, Halchenko YO, Waskom ML, Ghosh S. Nipype: a flexible, lightweight and extensible neuroimaging data processing framework in Python. Frontiers in Neuroinfor-matics. 2011; 5:13. doi: 10.3389/fninf.2011.00013.

Grady CL, Van Meter JW, Maisog JM, Pietrini P, Krasuski J, Rauschecker JP. Attention-related modulation of activity in primary and secondary auditory cortex. Neuroreport. 1997; 8(11):2511–2516.

Greve DN, Fischl B. Accurate and robust brain image alignment using boundary-based registration. NeuroImage. 2009; 48(1):63–72. doi: 10.1016/j.neuroimage.2009.06.060.

Grothe B, Pecka M, McAlpine D. Mechanisms of sound localization in mammals. Physiological reviews. 2010; 90(3):983–1012.

Gutschalk A, Patterson RD, Rupp A, Uppenkamp S, Scherg M. Sustained magnetic fields reveal separate sites for sound level and temporal regularity in human auditory cortex. Neuroimage. 2002; 15(1):207–216.

Hart HC, Hall DA, Palmer AR. The sound-level-dependent growth in the extent of fMRI activation in Heschl’s gyrus is different for low-and high-frequency tones. Hearing research. 2003; 179(1-2):104–112.

Hart HC, Palmer AR, Hall DA. Heschl’s gyrus is more sensitive to tone level than non-primary auditory cortex. Hearing research. 2002; 171(1-2):177–190.

Hartmann WM. Localization of sound in rooms. The Journal of the Acoustical Society of America. 1983; 74(5):1380–1391.

Hassager HG, Gran F, Dau T. The role of spectral detail in the binaural transfer function on perceived externalization in a reverberant environment. The Journal of the Acoustical Society of America. 2016; 139(5):2992–3000.

Haynes JD. Decoding mental states from brain activity in humans. Nature Reviews Neuroscience. 2006; 7(7):523–534. doi: 10.1038/nrn1931.

Hegerl U, Gallinat J, Mrowinski D. Intensity dependence of auditory evoked dipole source activity. International Journal of Psychophysiology. 1994; 17(1):1–13.

Hidaka T, Beranek LL, Okano T. Interaural cross-correlation, lateral fraction, and low-and high-frequency sound levels as measures of acoustical quality in concert halls. The Journal of the Acoustical Society of America. 1995; 98(2):988–1007.

Hoppe H, van de Par S, Flanagin V, Ewert SD. Combined assessment of auditory distance perception and externalization. arXiv [Preprint] (2024) https://arxivorg/abs/240814198. accessed 27 September 2024;.

ISO Central Secretary. Acoustics — Measurement of room acoustic parameters. Geneva, CH: International Organization for Standardization; 2009.

Jancke L, Shah N. Does dichotic listening probe temporal lobe functions? Neurology. 2002; 58(5):736–743.

Jäncke L, Shah N, Posse S, Grosse-Ryuken M, Müller-Gärtner HW. Intensity coding of auditory stimuli: an fMRI study. Neuropsychologia. 1998; 36(9):875–883.

Jäncke L, Mirzazade S, Shah NJ. Attention modulates activity in the primary and the secondary auditory cortex: a functional magnetic resonance imaging study in human subjects. Neuroscience letters. 1999; 266(2):125–128.

Jeffress LA. A place theory of sound localization. Journal of comparative and physiological psychology. 1948; 41(1):35.

Jenkinson M, Bannister P, Brady M, Smith S. Improved Optimization for the Robust and Accurate Linear Registration and Motion Correction of Brain Images. NeuroImage. 2002; 17(2):825–841. http://www.sciencedirect.com/science/article/pii/S1053811902911328, doi: 10.1006/nimg.2002.1132.

Kanwisher N. Functional specificity in the human brain: a window into the functional architecture of the mind. Proceedings of the national academy of sciences. 2010; 107(25):11163–11170.

Kell AJ, McDermott JH. Invariance to background noise as a signature of non-primary auditory cortex. Nature communications. 2019; 10(1):3958.

Kim HY, Suzuki Y, Takane S, Sone T. Control of auditory distance perception based on the auditory parallax model. Applied Acoustics. 2001; 62(3):245–270.

Klein A, Ghosh SS, Bao FS, Giard J, Häme Y, Stavsky E, Lee N, Rossa B, Reuter M, Neto EC, Keshavan A. Mindboggling morphometry of human brains. PLOS Computational Biology. 2017; 13(2):e1005350. http://journals.plos.org/ploscompbiol/article?id=10.1371/journal.pcbi.1005350, doi: 10.1371/journal.pcbi.1005350.

Klein F, Werner S, Mayenfels T. Influences of training on externalization of binaural synthesis in situations of room divergence. Journal of the Audio Engineering Society. 2017; 65(3):178–187.

Kolarik A, Cirstea S, Pardhan S. Discrimination of virtual auditory distance using level and direct-to-reverberant ratio cues. The Journal of the Acoustical Society of America. 2013 11; 134(5):3395–3398. 10.1121/1.4824395, doi: 10.1121/1.4824395.

Kolarik AJ, Moore BC, Zahorik P, Cirstea S, Pardhan S. Auditory distance perception in humans: a review of cues, development, neuronal bases, and effects of sensory loss. Attention, Perception, and Psychophysics. 2016; 78:373–395.

Kopčo N, Doreswamy KK, Huang S, Rossi S, Ahveninen J. Cortical auditory distance representation based on direct-to-reverberant energy ratio. NeuroImage. 2020; 208:116436. https://www.sciencedirect.com/science/article/pii/S1053811919310274, doi: 10.1016/j.neuroimage.2019.116436.

Kopčo N, Schoolmaster M, Shinn-Cunningham B. Learning to judge distance of nearby sounds in reverberant and anechoic environments. In: Proceedings of the Joint Congress CFA, DAGA ‘04 March 22-25, 2004, Strasbourg, France, vol. 4; 2004. p. 207–208.

Kopčo N, Shinn-Cunningham BG. Effect of stimulus spectrum on distance perception for nearby sources. The Journal of the Acoustical Society of America. 2011; 130(3):1530–1541.

Kopčo N, Huang S, Belliveau JW, Raij T, Tengshe C, Ahveninen J. Neuronal representations of distance in human auditory cortex. Proceedings of the National Academy of Sciences. 2012; 109(27):11019–11024. https://www.pnas.org/doi/abs/10.1073/pnas.1119496109, doi: 10.1073/pnas.1119496109.

Kriegeskorte N, Goebel R, Bandettini P. Information-based functional brain mapping. Proceedings of the National Academy of Sciences. 2006; 103(10):3863–3868.

Kriegeskorte N, Mur M, Bandettini P. Representational similarity analysis - connecting the branches of systems neuroscience. Frontiers in Systems Neuroscience. 2008; 2. https://www.frontiersin.org/articles/10.3389/neuro.06.004.2008, doi: 10.3389/neuro.06.004.2008.

Krumbholz K, Schönwiesner M, von Cramon DY, Rübsamen R, Shah NJ, Zilles K, Fink GR. Representation of interaural temporal information from left and right auditory space in the human planum temporale and inferior parietal lobe. Cerebral Cortex. 2005; 15(3):317–324.

Lanczos C. Evaluation of Noisy Data. Journal of the Society for Industrial and Applied Mathematics Series B Numerical Analysis. 1964; 1(1):76–85. http://epubs.siam.org/doi/10.1137/0701007, doi: 10.1137/0701007.

Langers DR, van Dijk P, Schoenmaker ES, Backes WH. fMRI activation in relation to sound intensity and loudness. Neuroimage. 2007; 35(2):709–718.

Larsen E, Iyer N, Lansing CR, Feng AS. On the minimum audible difference in direct-to-reverberant energy ratio. The Journal of the Acoustical Society of America. 2008 07; 124(1):450–461. 10.1121/1.2936368, doi: 10.1121/1.2936368.

Leclère T, Lavandier M, Perrin F. On the externalization of sound sources with headphones without reference to a real source. The Journal of the Acoustical Society of America. 2019; 146(4):2309–2320.

Lu Z, Ku Y. NeuroRA: A Python Toolbox of Representational Analysis From Multi-Modal Neural Data. Frontiers in neuroinformatics. 2020; 14.

MATLAB. version 9.13.0 (R2021b). Natick, Massachusetts, United States: The MathWorks Inc. 2017; https://www.mathworks.com.

McAlpine D. Creating a sense of auditory space. The Journal of Physiology. 2005; 566(1):21–28. doi: 10.1113/jphysiol.2005.083113.

McLaughlin SA, Higgins NC, Stecker GC. Tuning to binaural cues in human auditory cortex. Journal of the Association for Research in Otolaryngology. 2016; 17:37–53.

Mershon DH, Ballenger WL, Little AD, McMurtry PL, Buchanan JL. Effects of room reflectance and background noise on perceived auditory distance. Perception. 1989; 18(3):403–416.

Mershon DH, King LE. Intensity and reverberation as factors in the auditory perception of egocentric distance. Perception and Psychophysics. 1975; 18:409–415.

Moore BC, Glasberg BR. A revision of Zwicker’s loudness model. Acta Acustica united with Acustica. 1996; 82(2):335–345.

Mulert C, Jäger L, Propp S, Karch S, Störmann S, Pogarell O, Möller HJ, Juckel G, Hegerl U. Sound level dependence of the primary auditory cortex: Simultaneous measurement with 61-channel EEG and fMRI. Neuroimage. 2005; 28(1):49–58.

Neukirch M, Hegerl U, Kötitz R, Dorn H, Gallinat U, Herrmann W. Comparison of the amplitude/intensity function of the auditory evoked N1m and N1 components. Neuropsychobiology. 2002; 45(1):41–48.

Norbeck O, Sprenger T, Avventi E, Rydén H, Kits A, Berglund J, Skare S. Optimizing 3D EPI for rapid T1-weighted imaging. Magnetic Resonance in Medicine. 2020; 84(3):1441–1455. https://onlinelibrary.wiley.com/doi/abs/10.1002/mrm.28222, doi: 10.1002/mrm.28222.

Norman KA, Polyn SM, Detre GJ, Haxby JV. Beyond mind-reading: multi-voxel pattern analysis of fMRI data. Trends in Cognitive Sciences. 2006; 10(9):424–430. doi: 10.1016/j.tics.2006.07.005.

Palomäki KJ, Tiitinen H, Mäkinen V, May PJ, Alku P. Spatial processing in human auditory cortex: the effects of 3D, ITD, and ILD stimulation techniques. Cognitive brain research. 2005; 24(3):364–379.

Patriat R, Reynolds RC, Birn RM. An improved model of motion-related signal changes in fMRI. NeuroImage. 2017 Jan; 144, Part A:74–82. http://www.sciencedirect.com/science/article/pii/S1053811916304360, doi: 10.1016/j.neuroimage.2016.08.051.

Petkov CI, Kang X, Alho K, Bertrand O, Yund EW, Woods DL. Attentional modulation of human auditory cortex. Nature neuroscience. 2004; 7(6):658–663.

Philbeck JW, Mershon DH. Knowledge about typical source output influences perceived auditory distance. The Journal of the Acoustical Society of America. 2002 05; 111(5):1980–1983. 10.1121/1.1471899, doi: 10.1121/1.1471899.

Phillips D, Irvine D. Responses of single neurons in physiologically defined area AI of cat cerebral cortex: sensitivity to interaural intensity differences. Hearing research. 1981; 4(3-4):299–307.

Power JD, Mitra A, Laumann TO, Snyder AZ, Schlaggar BL, Petersen SE. Methods to detect, characterize, and remove motion artifact in resting state fMRI. NeuroImage. 2014; 84(Supplement C):320–341. http://www.sciencedirect.com/science/article/pii/S1053811913009117, doi: 10.1016/j.neuroimage.2013.08.048.

Quadflieg S, Mohr A, Mentzel HJ, Miltner WHR, Straube T. Modulation of the neural network involved in the processing of anger prosody: The role of task-relevance and social phobia. Biological Psychology. 2008; 78(2):129–137. doi: 10.1016/j.biopsycho.2008.01.014.

R Core Team. R: A Language and Environment for Statistical Computing. R Foundation for Statistical Computing, Vienna, Austria; 2023, https://www.R-project.org/.

Rämä P, Courtney SM. Functional topography of working memory for face or voice identity. Neuroimage. 2005; 24(1):224–234.

Rapin I, Schimmel H, Tourk LM, Krasnegor NA, Pollak C. Evoked responses to clicks and tones of varying intensity in waking adults. Electroencephalography and clinical neurophysiology. 1966; 21(4):335–344.

Rauschecker JP. Cortical processing of complex sounds. Current opinion in neurobiology. 1998; 8(4):516–521.

Rauschecker JP, Tian B. Mechanisms and streams for processing of “what” and “where” in auditory cortex. Proceedings of the National Academy of Sciences. 2000; 97(22):11800–11806.

Rauschecker JP, Tian B, Hauser M. Processing of complex sounds in the macaque nonprimary auditory cortex. Science. 1995; 268(5207):111–114.

Rauschecker J. Processing of complex sounds in the auditory cortex of cat, monkey, and man. Acta OtoLaryngologica. 1997; 117(sup532):34–38.

Reinhart P, Souza PE, Srinivasan NK, Gallun FJ. Effects of reverberation and compression on consonant identification in individuals with hearing impairment. Ear and Amp; Hearing. 2016; 37:144–152. doi: 10.1097/aud.0000000000000229.

Röhl M, Uppenkamp S. Neural coding of sound intensity and loudness in the human auditory system. Journal of the Association for Research in Otolaryngology. 2012; 13:369–379.

Satterthwaite TD, Elliott MA, Gerraty RT, Ruparel K, Loughead J, Calkins ME, Eickhoff SB, Hakonarson H, Gur RC, Gur RE, Wolf DH. An improved framework for confound regression and filtering for control of motion artifact in the preprocessing of resting-state functional connectivity data. NeuroImage. 2013; 64(1):240–256. http://linkinghub.elsevier.com/retrieve/pii/S1053811912008609, doi: 10.1016/j.neuroimage.2012.08.052.

Schroeder MR. New method of measuring reverberation time. The Journal of the Acoustical Society of America. 1965; 37(6_Supplement):1187–1188.

Schutte M, Aspects of room acoustics, vision and motion in the human auditory perception of space. Ludwig-Maximilians-Universität München; 2021. http://nbn-resolving.de/urn:nbn:de:bvb:19-285435.

Schwarzbauer C, Davis MH, Rodd JM, Johnsrude I. Interleaved silent steady state (ISSS) imaging: A new sparse imaging method applied to auditory fMRI. NeuroImage. 2006; 29(3):774–782. https://www.sciencedirect.com/science/article/pii/S1053811905006063, doi: 10.1016/j.neuroimage.2005.08.025.

Shinn-Cunningham B. Learning reverberation: Considerations for spatial auditory displays. Proceedings of the 2000 International Conference on Auditory Display. 2000;.

Somers DC, Dale AM, Seiffert AE, Tootell RB. Functional MRI reveals spatially specific attentional modulation in human primary visual cortex. Proceedings of the National Academy of Sciences. 1999; 96(4):1663–1668.

Stecker GC, McLaughlin SA, Higgins NC. Monaural and binaural contributions to interaural-level-difference sensitivity in human auditory cortex. Neuroimage. 2015; 120:456–466.

Stevens KN. In: Rigault A, Charbonneau R, editors. Sources of Inter- and Intra-Speaker Variability in the Acoustic Properties of Speech Sounds Berlin, Boston: De Gruyter Mouton; 1972. p. 206–232. 10.1515/9783110814750-014, doi: doi:10.1515/9783110814750-014.

Stirnberg R, Huijbers W, Brenner D, Poser BA, Breteler M, Stöcker T. Rapid Whole-Brain Resting-State fMRI at 3 Tesla: Efficiency-optimized Three-Dimensional EPI versus Repetition Time-Matched Simultaneous-Multi-Slice EPI. NeuroImage. 2017; 163(August):81–92. http://linkinghub.elsevier.com/retrieve/pii/S105381191730678X, doi: 10.1016/j.neuroimage.2017.08.031.

Stirnberg R, Stöcker T. Segmented K-space blipped-controlled aliasing in parallel imaging for high spatiotem-poral resolution EPI. Magnetic resonance in medicine. 2021; 85(3):1540–1551.

Sutojo S, Thiemann J, Kohlrausch A, van de Par S. In: Blauert J, Braasch J, editors. Auditory Gestalt Rules and Their Application Cham: Springer International Publishing; 2020. p. 33–59. 10.1007/978-3-030-00386-9_2, doi: 10.1007/978-3-030-00386-9_2.

Tata MS, Ward LM. Early phase of spatial mismatch negativity is localized to a posterior “where” auditory pathway. Experimental Brain Research. 2005; 167:481–486.

Thompson GC, Cortez AM. The inability of squirrel monkeys to localize sound after unilateral ablation of auditory cortex. Behavioural brain research. 1983; 8(2):211–216.

Toole FE. In-Head Localization of Acoustic Images. The Journal of the Acoustical Society of America. 1970; 48(4B):943–949.

Tustison NJ, Avants BB, Cook PA, Zheng Y, Egan A, Yushkevich PA, Gee JC. N4ITK: Improved N3 Bias Correction. IEEE Transactions on Medical Imaging. 2010; 29(6):1310–1320. doi: 10.1109/TMI.2010.2046908.

Ungan P, Yagcioglu S, Goksoy C. Differences between the N1 waves of the responses to interaural time and intensity disparities: scalp topography and dipole sources. Clinical Neurophysiology. 2001; 112(3):485–498.

Von Békésy G. Über die Entstehung der Entfernungsempfindung beim Hören. Akust Z. 1938; 3:21–31.

Warren JD, Zielinski BA, Green GG, Rauschecker JP, Griffiths TD. Perception of sound-source motion by the human brain. Neuron. 2002; 34(1):139–148.

Wendt T, van de Par S, Ewert SD. A Computationally-Efficient and Perceptually-Plausible Algorithm for Binaural Room Impulse Response Simulation. J Audio Eng Soc. 2014; 62(11):748–766. https://www.aes.org/e-lib/browse.cfm?elib=17550.

Werner S, Klein F, Mayenfels T, Brandenburg K. A summary on acoustic room divergence and its effect on externalization of auditory events. In: 2016 Eighth International Conference on Quality of Multimedia Experience (QoMEX); 2016. p. 1–6. doi: 10.1109/QoMEX.2016.7498973.

Wessinger CM, VanMeter J, Tian B, Van Lare J, Pekar J, Rauschecker JP. Hierarchical organization of the human auditory cortex revealed by functional magnetic resonance imaging. Journal of cognitive neuroscience. 2001; 13(1):1–7.

Wisniewski MG, Mercado III E, Gramann K, Makeig S. Familiarity with speech affects cortical processing of auditory distance cues and increases acuity. PLoS One. 2012; 7(7):e41025.

Xu J, Dong H, Li N, Wang Z, Guo F, Wei J, Dang J. Weighted RSA: An Improved Framework on the Perception of Audio-visual Affective Speech in Left Insula and Superior Temporal Gyrus. Neuroscience. 2021; 469:46–58. https://www.sciencedirect.com/science/article/pii/S0306452221002876, doi: 10.1016/j.neuroscience.2021.06.002.

Yin TCT, Smith PH, Joris PX, Neural mechanisms of binaural processing in the auditory brainstem. Wiley; 2019.

You J, Jeon JY. Just noticeable differences in sound quality metrics for refrigerator noise. Noise Control Engineering Journal - NOISE CONTR ENG J. 2008 11; 56. doi: 10.3397/1.2987734.

Zahorik P. Assessing auditory distance perception using virtual acoustics. The Journal of the Acoustical Society of America. 2002; 111(4):1832–1846.

Zahorik P. Direct-to-reverberant energy ratio sensitivity. The Journal of the Acoustical Society of America. 2002 10; 112(5):2110–2117. 10.1121/1.1506692, doi: 10.1121/1.1506692.

Zahorik P, Brungart DS, Bronkhorst AW. Auditory distance perception in humans: A summary of past and present research. ACTA Acustica united with Acustica. 2005; 91(3):409–420.

Zatorre RJ, Belin P. Spectral and temporal processing in human auditory cortex. Cerebral cortex. 2001; 11(10):946–953.

Zatorre RJ, Penhune VB. Spatial localization after excision of human auditory cortex. Journal of Neuroscience. 2001; 21(16):6321–6328.

Zhang J, Nakamoto KT, Kitzes LM. Binaural interaction revisited in the cat primary auditory cortex. Journal of neurophysiology. 2004; 91(1):101–117.

Zhang Y, Brady M, Smith S. Segmentation of brain MR images through a hidden Markov random field model and the expectation-maximization algorithm. IEEE Transactions on Medical Imaging. 2001; 20(1):45–57. doi: 10.1109/42.906424.

